# Disease-specific variant pathogenicity prediction significantly improves variant interpretation in inherited cardiac conditions

**DOI:** 10.1101/2020.03.27.010736

**Authors:** Xiaolei Zhang, Roddy Walsh, Nicola Whiffin, Rachel Buchan, William Midwinter, Alicja Wilk, Risha Govind, Nicholas Li, Mian Ahmad, Francesco Mazzarotto, Angharad Roberts, Pantazis Theotokis, Erica Mazaika, Mona Allouba, Antonio de Marvao, Chee Jian Pua, Sharlene M Day, Euan Ashley, Steven D Colan, Michelle Michels, Alexandre C Pereira, Daniel Jacoby, Carolyn Y Ho, Iacopo Olivotto, Gunnar T Gunnarsson, John Jefferies, Chris Semsarian, Jodie Ingles, Declan P. O’Regan, Yasmine Aguib, Magdi H. Yacoub, Stuart A. Cook, Paul J.R. Barton, Leonardo Bottolo, James S. Ware

**Affiliations:** National Heart and Lung Institute, Imperial College London, London, United Kingdom; Cardiovascular Research Centre, Royal Brompton and Harefield NHS, Foundation Trust London, London, United Kingdom; Oxford Medical Genetics Laboratory, Oxford University Hospitals NHS Foundation Trust, The Churchill Hospital, Oxford, United Kingdom; Radcliffe Department of Medicine, University of Oxford, Oxford, United Kingdom; MRC London Institute of Medical Sciences, Imperial College London, London, United Kingdom; Cardiomyopathy Unit, Careggi University Hospital, Florence, Italy; Department of Clinical and Experimental Medicine, University of Florence, Florence, Italy; Aswan Heart Centre, Magdi Yacoub Heart Foundation, Aswan, Egypt; National Heart Centre Singapore, Singapore; Division of Cardiovascular Medicine and Penn Cardiovascular Institute, Perelman School of Medicine, University of Pennsylvania, Philadelphia, USA; Division of Cardiovascular Medicine, Stanford University Medical Center, Stanford, CA, USA; Department of Cardiology, Boston Children’s Hospital, Boston, MA, USA; Department of Cardiology, Thoraxcenter, Erasmus MC Rotterdam, The Netherlands; Heart Institute (InCor), University of Sao Paulo Medical School, Sao Paulo, Brazil; Yale University, New Haven, CT, USA; Cardiovascular Division, Brigham and Women’s Hospital, Boston, MA, USA; Faculty of Medicine, University of Iceland, Akureyri, Iceland; Cincinnati Children’s Hospital, University of Cincinnati Department of Pediatrics, Cincinnati, OH, USA; Centenary Institute, The University of Sydney, Sydney, Australia; Department of Cardiology, Royal Prince Alfred Hospital, Sydney, Australia; Duke-National University of Singapore, Singapore; Department of Medical Genetics, University of Cambridge, Cambridge, United Kingdom; Alan Turing Institute, London, United Kingdom; MRC Biostatistics Unit, University of Cambridge, Cambridge, United Kingdom

**Keywords:** pathogenicity prediction, variant interpretation, missense variant, cardiomyopathy, Long QT syndrome, Brugada syndrome

## Abstract

**Background:** Accurate discrimination of benign and pathogenic rare variation remains a priority for clinical genome interpretation. State-of-the-art machine learning tools are useful for genome-wide variant prioritisation but remain imprecise. Since the relationship between molecular consequence and likelihood of pathogenicity varies between genes with distinct molecular mechanisms, we hypothesised that a disease-specific classifier may outperform existing genome-wide tools.

**Methods:** We present a novel disease-specific variant classification tool, CardioBoost, that estimates the probability of pathogenicity for rare missense variants in inherited cardiomyopathies and arrhythmias, trained with variants of known clinical effect. To benchmark against state-of-the-art genome-wide pathogenicity classification tools, we assessed classification of hold-out test variants using both overall performance metrics, and metrics of high-confidence (>90%) classifications relevant to variant interpretation. We further evaluated the prioritisation of variants associated with disease and patient clinical outcomes, providing validations that are robust to potential mis-classification in gold-standard reference datasets.

**Results:** CardioBoost has higher discriminating power than published genome-wide variant classification tools in distinguishing between pathogenic and benign variants based on overall classification performance measures with the highest area under the Precision-Recall Curve as 91% for cardiomyopathies and as 96% for inherited arrhythmias. When assessed at high-confidence (>90%) classification thresholds, prediction accuracy is improved by at least 120% over existing tools for both cardiomyopathies and arrhythmias, with significantly improved sensitivity and specificity. Finally, CardioBoost improves prioritisation of variants significantly associated with disease, and stratifies survival of patients with cardiomyopathies, confirming biologically relevant variant classification.

**Conclusions:** We demonstrate that a disease-specific variant pathogenicity prediction tool outperforms state-of-the-art genome-wide tools for the classification of rare missense variants of uncertain significance for inherited cardiac conditions. To facilitate evaluation of CardioBoost, we provide pre-computed pathogenicity scores for all possible rare missense variants in genes associated with cardiomyopathies and arrhythmias (https://www.cardiodb.org/cardioboost/). Our results also highlight the need to develop and evaluate variant classification tools focused on specific diseases and clinical application contexts. Our proposed model for assessing variants in known disease genes, and the use of application-specific evaluations, is broadly applicable to improve variant interpretation across a wide range of Mendelian diseases.

## Background

The accurate prediction of the effect of a previously unseen genetic variant on disease risk is an unmet need in clinical genetics. According to guidelines developed by the American College of Medical Genetics and Genomics/Association for Molecular Pathology (ACMG/AMP)^1^, computational prediction of variant pathogenicity is integrated as one line of supporting evidence to assess the clinical significance of human genetic variation. Several tools have been developed to predict the effects of rare variants given multiple functional annotations, such as evolutionary conservation scores and biochemical properties, and to derive scores describing the likelihood of pathogenicity^2–6^. Recent efforts have employed state-of-the-art machine learning classification methods including ensemble learning^7,8^ and deep learning^9^ to improve predictions.

While existing genome-wide variant classification tools learn from large-scale data over the entire genome, they might also compromise the prediction accuracy for specific sets of genes and diseases^10^ in the following ways. First, variation in a single gene can cause distinct clinical phenotypes via different allelic mechanisms. Genome-wide machine learning tools that classify variants as deleterious or not, without reference to a specific disease or mechanism, may not perform as well as those that separate gene-disease relations since, for example, they do not distinguish between gain- and loss-of-function variants. Second, genome-wide classification tools may not benefit from specific lines of evidence only available for a subset of well-characterised genes or diseases. We have previously shown^11^ that the addition of gene- and disease-specific evidence into a transparent Bayesian model improves variant interpretation in inherited cardiac diseases. Finally, most genome-wide prediction tools are reported to have low specificity^1^.

Furthermore, the measures used in the evaluation of existing machine learning variant classification tools are not always well defined or the most clinically-relevant. The performance of variant classification is routinely evaluated using conventional classification performance measures such as the receiver operating characteristic (ROC) curve, that assesses diagnostic performance across a range of discrimination thresholds, or metrics such as sensitivity and specificity derived from the confusion matrix at a single, specified threshold. We argue that these measures should be tailored to the specific application at hand. In particular, it is necessary to consider the relative cost of decisions based on the Type I and Type II errors in any specific application, as different contexts may favour the control of Type I error (limiting false positive assertions) or Type II error (limiting false negative assertions). For example, when classifying a variant for predictive genetic testing, control of the Type I error is usually prioritised: familial cascade testing on a variant falsely reported as pathogenic can be extremely harmful^12^. Conversely, if considering whether to offer a patient a therapy proven to be effective in a subgroup of patients with a particular molecular aetiology (e.g., Sulfonylureas in some types of monogenic diabetes^13^), one might prioritise the control of Type II error, since it is important to identify all who might benefit from targeted treatment when its benefits outweigh the side-effects. Most current variant classifier tools favour sensitivity over control of the Type I error with over-prediction of pathogenic variants^1^. The inappropriate use of performance measures not only affects the construction of the best classifier, but also the evaluation of its utility in clinical applications.

To address the disadvantages of using genome-wide classification tools, we sought to develop an accurate variant classifier considering gene-disease relations by taking inherited cardiac conditions (ICCs) as examples. The resulting disease-specific variant classification tool, CardioBoost, includes two disease-specific variant classifiers for two groups of closely related syndromes: one classifier for familial cardiomyopathies (CM) that include hypertrophic cardiomyopathy (HCM) and dilated cardiomyopathy (DCM), and the other for inherited arrythmia syndromes (IAS) that include long QT syndrome (LQTS) and Brugada syndrome.

While optimally it may be desirable to train a specific model for every gene-disease pair, this is not feasible due to current limitations in the number of variants with well-characterised disease consequences for training (and testing). Moreover we have previously demonstrated benefit from jointly-fitting some parameters across closely-related genes or diseases^11^.We therefore constructed models that aggregate related genes as described above, hypothesising that these disease-specific models are biologically plausible since the relevance of computational evidence types to interpret variant effect is more likely transferable within closely related syndromes.

Trained on well-curated disease-specific data, CardioBoost integrates multiple variant annotations and pathogenicity scores obtained from previously published computational tools, and predicts the probability that rare missense variants are pathogenic for monogenic inherited cardiac conditions, based on the Adaptive Boosting (AdaBoost) algorithm^13^. Our tool has improved performances over state-of-the-art genome-wide tools in a variety of tasks including separation of pathogenic from benign variants and prioritisation of variants highly associated with disease and adverse clinical outcomes.

## Methods

### Building CardioBoost

A full description of data collection, model development and validation is given in the **Supplementary Methods**. In brief, we constructed two classifiers, one for inherited cardiomyopathies, and one for inherited arrhythmia syndromes, to output the estimated probability of pathogenicity for rare missense variants in genes robustly associated with these diseases. The CM classifier is applicable for 16 genes associated with hypertrophic and dilated cardiomyopathies. To obtain training and test sets, we collected 356 unique rare (gnomAD minor allele frequency < 0.1%) missense variants in established cardiomyopathy-associated genes (**Supplementary Table 1**) identified in 9,007 individuals either with a confirmed clinical diagnosis of CM, or referred for genetic testing with a diagnosis of CM, and interpreted as Pathogenic or Likely Pathogenic. For the inherited arrhythmia classifier, we consider genes associated with long QT syndrome and Brugada syndrome. 252 unique rare missense variants reported to be Pathogenic or Likely Pathogenic with no conflicting interpretations (Benign or Likely benign) in established arrhythmia-associated genes (**Supplementary Table 2**) were collected from NCBI ClinVar Database^14^. As a benign variant set, 302 unique rare missense variants in cardiomyopathy genes, and 237 unique rare missense variants in arrhythmia genes were collected from the targeted sequencing of 2,090 healthy volunteers. Since these volunteers have no family history of ICCs and confirmed without ICCs on ECG or cardiac MRI, this cohort provides a lower disease prevalence than a general population thus the rare missense variants carried by them shall be considered as highly likely benign to inherited cardiac conditions. To avoid over-fitting, for each condition the data set were randomly split, with two-thirds used for training and one-third reserved as a hold-out test set (**Supplementary Table 3-5**).

For each variant, we collected 76 functional annotations (**Supplementary Table 6** and **Supplementary Methods**) as features in our disease-specific variant classification tool, including intra- and inter-species conservation scores, amino acid substitution scores, and pathogenicity predictions from published genome-wide variant classifiers. We selected nine classification algorithms including best-in-class representatives of all of the major families of machine learning algorithms, and applied a nested cross-validation^15^ to select the optimal algorithm for our tool. In the inner 5-fold cross-validation loop, a candidate classification algorithm was trained in order to optimise its hyper-parameters. In the outer 10-fold cross-validation loop, the optimised candidate algorithms were compared and the best-performing one was selected (see **Figure 1** and **Supplementary Methods**).

**Figure 1.**
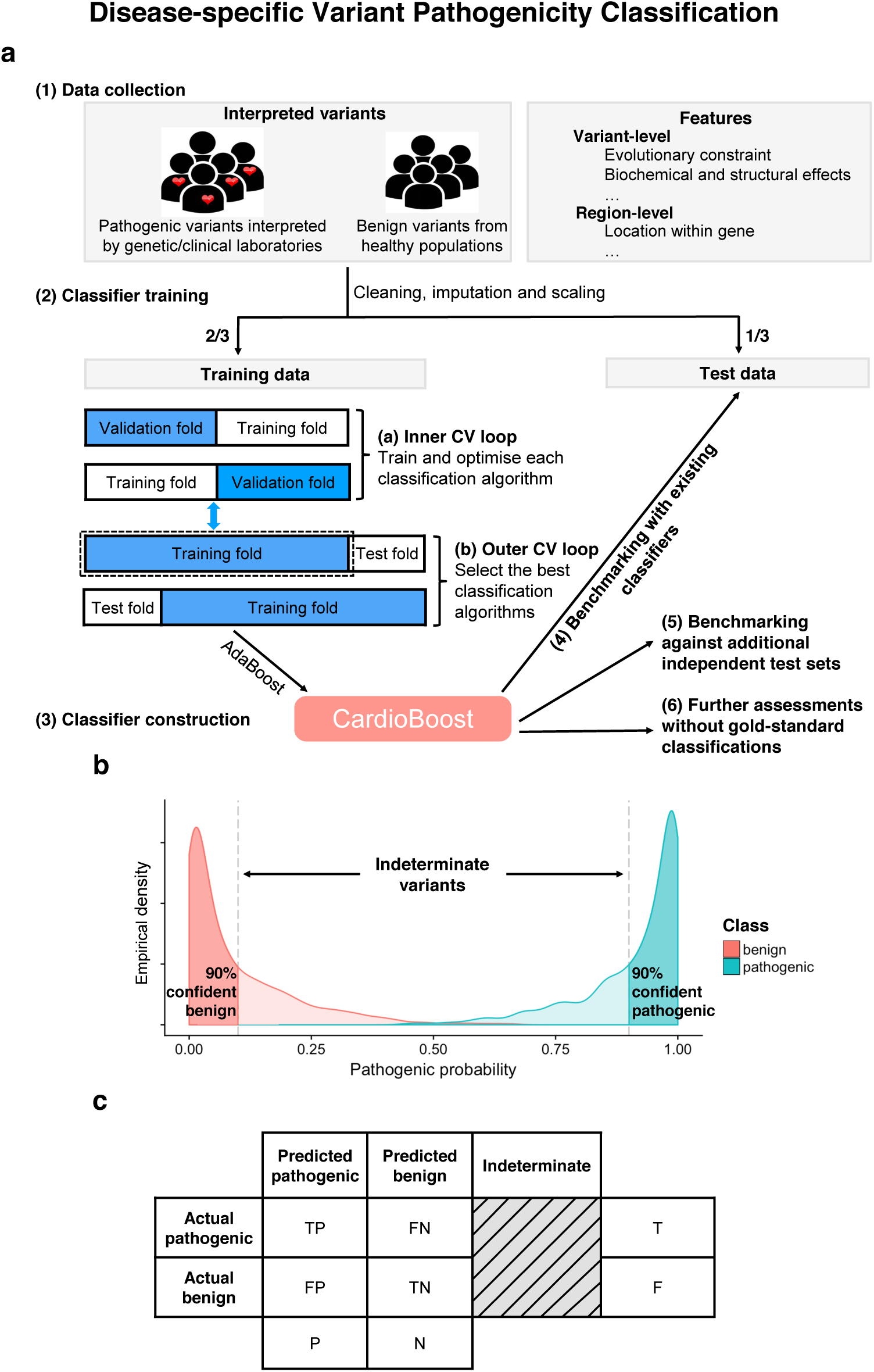
Training, and testing of CardioBoost, and definition of high-confidence variant classification thresholds for performance assessment. (**a**) Construction of CardioBoost: (1) After defining gold-standard data, (2) the dataset was split with a 2:1 proportion into training and test tests. The training set was used for two rounds of cross-validation: first to optimise individually a number of possible machine learning algorithms, and second to select the best performing tool. (3) AdaBoost was the best performing algorithm, and forms the basis of CardioBoost. (4) CardioBoost was benchmarked against existing best-in-class tools using the hold-out test data, (5) a number of additional independent test sets, and (6) approaches based on association with clinical characteristics of variant carriers that do not rely on a gold-standard classification. (**b**) Illustrative distributions of predicted pathogenicity scores for a set of pathogenic and benign variants obtained by a hypothetical binary classifier. In a clinical context (based on ACMG/AMP guidelines), variants are classified into the following categories according to the probability of pathogenicity: Pathogenic/Likely Pathogenic (Probability of pathogenicity (Pr) >=0.9), Benign/Likely Benign (Pr <=0.1) and a clinically indeterminate group of Variants of Uncertain Significance with low interpretative confidence (0.1 < Pr < 0.9). (**c**)The corresponding confusion matrix with the defined double classification thresholds Pr >=0.9 and Pr <=0.1.

For both conditions, AdaBoost^13^ was selected with the best cross-validated out-of-sample performance (see **Supplementary Methods** and **Supplementary Table 7-8**). AdaBoost is a boosting tree classification algorithm combining many decision trees. Each decision tree is learned sequentially to assign more weight to samples misclassified by the previous decision tree, and weighted by its classification accuracy. Having selected AdaBoost as the basis for our disease-specific classifier, a predictive model was constructed by training AdaBoost on the whole training set, to produce a final variant classification model for each disease, named CardioBoost.

CardioBoost was benchmarked against genome-wide classification tools using an unseen hold-out test set. We applied conventional global classification performance measures, as well as specific measures focusing on high-confidence thresholds. To ensure robustness, we further assessed for prioritisation of variants associated with disease in independent cohorts and associated with patients’ survival measures. These two approaches are relatively independent of the gold-standard classification from human experts’ interpretation, and directly assess the relationship between the clinical phenotype and the prioritised variants (for the descriptions of the benchmarking methods see **Supplementary Methods**).

## Results

### CardioBoost outperforms state-of-the-art genome-wide prediction tools based on conventional classification performance measures

The hold-out test sets were used to evaluate the classifiers’ performance on unseen data. CardioBoost was compared against two recently developed genome-wide variant classification algorithms, M-CAP and REVEL, reported to have leading performance in pathogenicity prediction of rare missense variants. Classification performance was first summarised using the area under the Precision-Recall Curve^16^ (PR-AUC), the area under the Receiver Operating Characteristic Curve (ROC-AUC) and Brier Score^17^, without relying on a single pre-defined classification threshold to discriminate pathogenic and benign variants.

In both inherited cardiac conditions, CardioBoost achieved the best values in all the three measures (**Figure 2**). The difference in performance was statistically significant for cardiomyopathies, with significantly increased PR-AUC (maximum *P*-value = 0.005 between the pairwise statistical comparisons of CardioBoost vs. M-CAP and CardioBoost vs. REVEL via permutation test), ROC-AUC (maximum *P*-value = 5×10^−6^ between the pairwise statistical comparisons using Delong test^18^), and Brier Score (maximum *P*-value = 0.005 between the pairwise comparisons via permutation test). CardioBoost also has significantly improved the Brier Score for arrhythmia syndromes (maximum *P*-value = 0.02 between the pairwise comparisons via permutation test).

**Figure 2.**
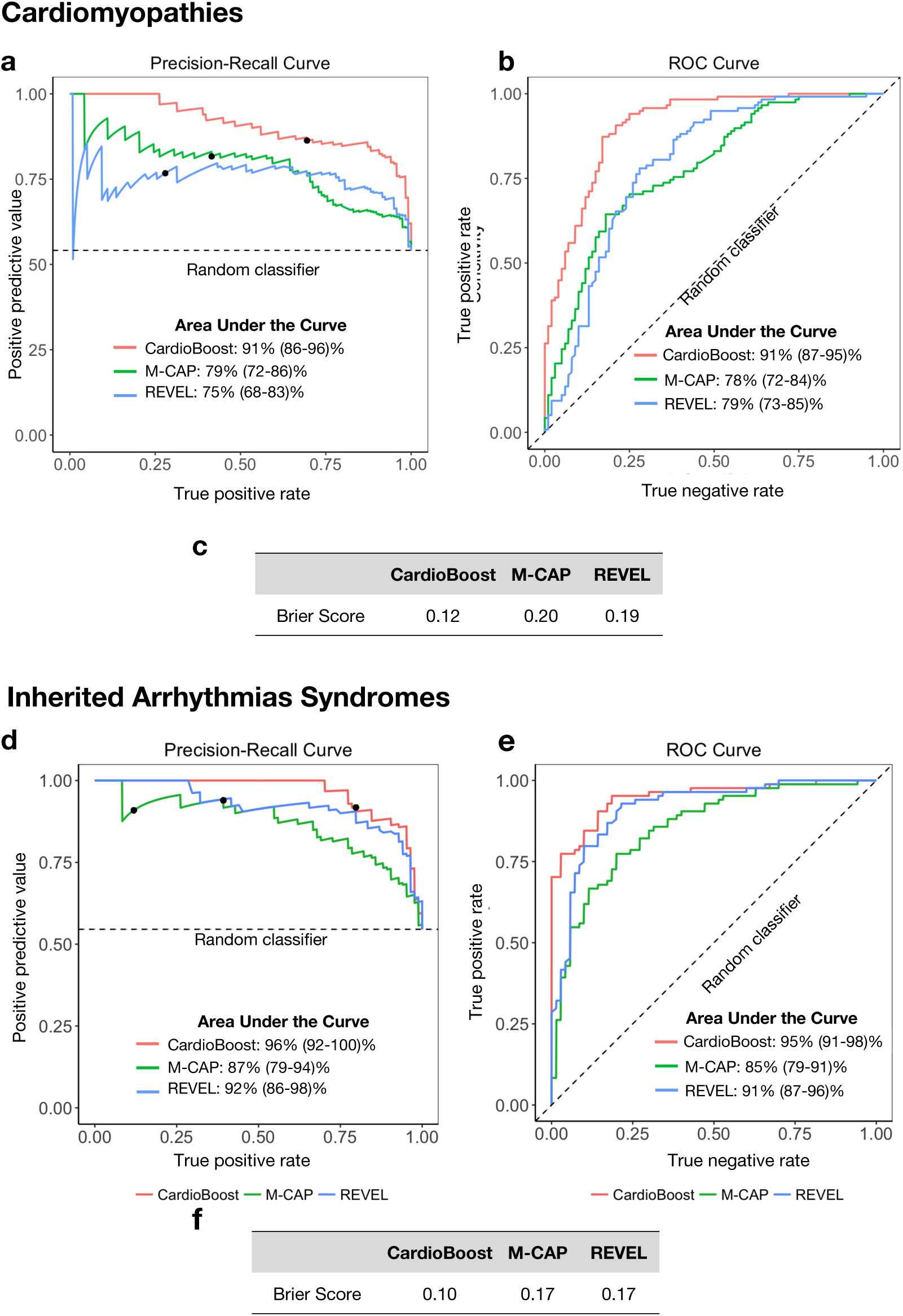
CardioBoost outperforms genome-wide prediction tools on hold-out test data. (**a**-**c**) Precision-Recall Curves, ROC Curves and Brier Scores for cardiomyopathy variant pathogenicity prediction. (**d**-**f**) Precision-Recall Curves, ROC Curves and Brier Score for inherited arrhythmia variant pathogenicity prediction. In (**a**) and (**d**), the marked point (•) indicates the precision (positive predictive value) and recall (true positive rate) at the 90% confidence level defined as clinically reportable in international guidelines. The dashed lines demonstrate the performance of a random classifier.

While CardioBoost was trained and tested on independent datasets, some variants had been used previously in the training of M-CAP and REVEL, whose pathogenicity scores were used as input features for CardioBoost (**Supplementary Table 6**). Thus, CardioBoost has been indirectly exposed to these variants. This may worsen classification performance if the variants were erroneously labelled during upstream training, or lead to artificially inflated performance estimates through concealed overfitting. To estimate the extent to which these potential limitations affect the prediction performance, we performed a stratification analysis to compare the performance of CardioBoost on variants used to train upstream genome-wide learners (indirectly “seen”), and variants that were completely novel (“unseen”) in the hold-out test data set. CardioBoost improved on cardiomyopathy- and arrhythmia-specific prediction over existing genome-wide classification tools both on indirectly “seen” (used in the training of M-CAP and REVEL) and “unseen” data. The overall accuracy of CardioBoost between the unseen and seen data sets is not significantly different for either CM or IAS. (**Supplementary Table 9-10** and **Supplementary Methods**).

### CardioBoost outperforms existing genome-wide prediction tools on high-confidence classification measures

In addition to estimating conventional classification performance, we evaluated performance at thresholds corresponding to accepted levels of certainty required for clinical decision making^1^ (90%; see definitions on **Figure 1b, Figure 1c** and **Supplementary Methods**). Using these thresholds (Pathogenic/Likely Pathogenic: probability of pathogenicity (Pr) ≥ 0.9; Benign/Likely Benign: Pr ≤ 0.1; Indeterminate: 0.1 < Pr < 0.9), CardioBoost again outperforms existing genome-wide machine learning variant classification tools when assessed using hold-out test data (**Table 1**).

**Table 1.**
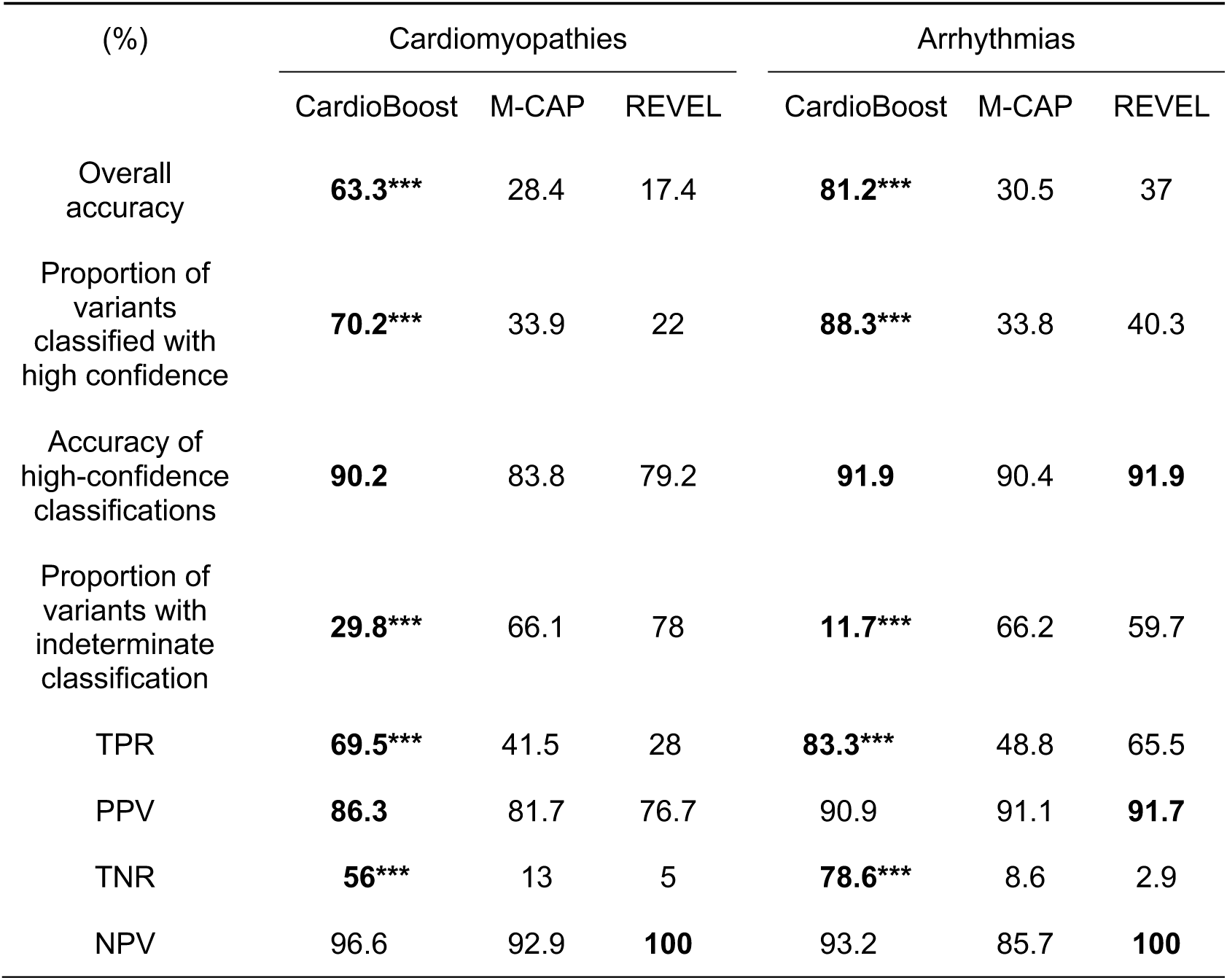
CardioBoost outperforms existing genome-wide tools for the classification of hold-out test variants. The performance of each tool is reported using the clinically relevant variant classification thresholds: high-confidence pathogenic (Pr ≥ 0.9), high-confidence benign (Pr ≤ 0.1), and indeterminate. For each predictive performance measure (see **Supplementary Methods** for details) the best algorithm is highlighted in bold. Permutation tests were performed to evaluate whether the performance of CardioBoost was significantly different from the best value obtained by M-CAP or REVEL (significance levels: \*\*\**P*-value ≤ 0.001, \*\**P*-value ≤ 0.01, \**P*-value ≤ 0.05).

| (%) | Cardiomyopathies |  |  | Arrhythmias |  |  |
| --- | --- | --- | --- | --- | --- | --- |
|  | CardioBoost | M-CAP | REVEL | CardioBoost | M-CAP | REVEL |
| Overall accuracy | **63.3\*\*\*** | 28.4 | 17.4 | **81.2\*\*\*** | 30.5 | 37 |
| Proportion of variants classified with high confidence | **70.2\*\*\*** | 33.9 | 22 | **88.3\*\*\*** | 33.8 | 40.3 |
| Accuracy of high-confidence classifications | **90.2** | 83.8 | 79.2 | **91.9** | 90.4 | **91.9** |
| Proportion of variants with indeterminate classification | **29.8\*\*\*** | 66.1 | 78 | **11.7\*\*\*** | 66.2 | 59.7 |
| TPR | **69.5\*\*\*** | 41.5 | 28 | **83.3\*\*\*** | 48.8 | 65.5 |
| PPV | **86.3** | 81.7 | 76.7 | 90.9 | 91.1 | **91.7** |
| TNR | **56\*\*\*** | 13 | 5 | **78.6\*\*\*** | 8.6 | 2.9 |
| NPV | 96.6 | 92.9 | **100** | 93.2 | 85.7 | **100** |

CardioBoost also maximises the identification of both pathogenic and benign variants. In both conditions, the proposed variant classification model had the highest true positive rate (TPR) (CM 69.5%; IAS 83.3%) and true negative rate (TNR) (CM 56%; IAS 78.6%) (*P*-value < 0.001). In total, CardioBoost correctly classified 63.3% of cardiomyopathy test variants and 81.2% of arrhythmia test variants with 90% or greater confidence-level. Such proportions of correctly classified variants are significantly higher (*P*-value < 0.001) than those obtained with M-CAP (CM 28.4%; IAS 30.5%) and REVEL (CM 17.4%; IAS 37%). In addition, CardioBoost minimises the number of indeterminate variants. Only 29.8% of cardiomyopathy test variants and 11.7% of arrhythmia test variants achieved indeterminate scores between 0.1 and 0.9, which were significantly fewer (*P*-value < 0.001) than those obtained with M-CAP (CM 66.1%; IAS 66.2%) or REVEL (CM 78%; IAS 59.7%) (**Table 1**).

Overall, using these thresholds CardioBoost assigned high-confidence classifications to 70.2% of cardiomyopathy test variants, among which 90.2% were correct. For arrhythmias, CardioBoost reported 88.3% of test variants with high confidence, with 91.9% prediction accuracy. The reported results are robust to the choice of classification thresholds. While guidelines propose 90% confidence as appropriate thresholds for likely pathogenic or likely benign classifications, some may advocate a higher confidence threshold. When assessed at a 95%-certainty classification threshold, CardioBoost continues to consistently outperform genome-wide tools with significantly (*P*-value < 0.001) higher accuracies (**Supplementary Table 11**).

CardioBoost is not intended to replace a full expert variant assessment in clinical practice, but for comparative purposes it is informative to consider how classification performance changes under application in different contexts. PPV and NPV are both dependent on the proportion of pathogenic variants in the variant set being tested, and so it is important to consider how our benchmarking translates to real-world application. Here we used the TPR and TNR calculated on our hold-out benchmarking test set to derive estimates of PPV and NPV for CardioBoost applied in different contexts where the true proportion of pathogenic variants might differ. Our estimation provides a lower bound of PPV and NPV under the assumption that pathogenic variants are fully penetrant. In the context of predictive genetic testing, the limitation of false positive prediction is prioritised, necessitating conservative estimates of PPV. Here we estimate reasonably conservative PPVs and corresponding NPVs of CardioBoost applied in two scenarios: in a diagnostic referral series and in samples from a general population. In a diagnostic laboratory cardiomyopathy referral series, where we estimate approximately 60% rare missense variants found in cardiomyopathy-associated genes to be pathogenic, the PPV and NPV of CardioBoost were estimated at 89% and 96% respectively. By contrast, if applied to variants in the same genes in a general population, where we estimate the proportion of rare variants that are pathogenic as ∼ 1%, the PPV and NPV reach 5% and 99.9%. Similarly, we estimated the performance of CardioBoost in an arrhythmia cohort (PPV: 95%; NPV: 87%) and a general population (PPV:3%; NPV: 99.9%). This suggests that the predictions of pathogenicity by CardioBoost are calibrated for high confidence only when applied in a diagnostic context, as would be expected. Classifications are appropriate for variants found in individuals with disease, with a reasonable prior probability of pathogenicity (the estimation details are described in **Supplementary Methods**).

Finally, as novel pathogenic variants are more likely to be ultra-rare (Minor allele frequency < 0.01%), we also tested CardioBoost performance on a hold-out set of only ultra-rare variants and confirmed that it consistently outperforms existing genome-wide tools (**Supplementary Table 12**). Its performance on ultra-rare variants is comparable with that on rare variants.

### Replication on additional independent test data confirms that CardioBoost improves prediction of pathogenic and benign variants

We collected four additional sets of independent test data to further assess the CardioBoost performance, using variants reported as pathogenic in ClinVar and HGMD^19^ (both databases of aggregated classified variants), a diagnostic laboratory referral series from the Oxford Molecular Genetics Laboratory (OMGL), and a large registry of HCM patients, SHaRe^20^. CardioBoost consistently achieved the highest TPRs: predicting the most pathogenic variants with over 90% certainty (**Table 2**). On a set of rare variants found in the gnomAD reference dataset, which is not enriched for inherited cardiac conditions and hence where the prevalence of disease should be equivalent to the general population, CardioBoost consistently predicts the most variants as benign (**Table 2**). CardioBoost also performed best when assessed at a higher 95%-certainty classification threshold (**Supplementary Table 13**) and on sets of ultra-rare variants (**Supplementary Table 14**).

**Table 2.** Evaluation of performances on additional test sets. CardioBoost performance was evaluated against additional variant sets. Four resources provided known pathogenic variants (SHaRe cardiomyopathy registry, ClinVar (two-star submissions), a UK regional genetic laboratory (Oxford Medical Genetics Laboratory – OMGL) and the Human Gene Mutation Database – HGMD). Variants found in gnomAD population controls were expected to be predominantly benign. Since gnomAD includes variants seen in the previous ExAC dataset that was partially used to train M-CAP and REVEL, we tested against the subset of variants in gnomAD that were not in ExAC. The number of variants in each set is shown in brackets. The TPR is reported for pathogenic variant test sets (with threshold Pr ≥ 0.9), and the TNR for benign variant test sets (with threshold Pr ≤ 0.1). For each performance measure the best algorithm is highlighted in bold. Permutation tests were carried out to evaluate whether the performance of CardioBoost was significantly different from the best value obtained by M-CAP or REVEL (significance levels: \*\*\**P*-value ≤ 0.001, \*\**P*-value ≤ 0.01, \**P*-value ≤ 0.05)

|  | Cardiomyopathies |  |  |  |
| --- | --- | --- | --- | --- |
|  | Pathogenic test variants (TPR) |  |  | Benign/population test variants (TNR) |
|  | SHaRe (N = 129) | ClinVar (N = 15) | HGMD (N = 145) | gnomAD (N = 2,003) |
| CardioBoost | **62.0\*\*\*** | **66.7** | **41.4\*\*\*** | **51.5\*\*\*** |
| M-CAP | 37.2 | 40.0 | 22.1 | 20.3 |
| REVEL | 24.0 | 53.3 | 22.8 | 5.6 |
|  | Arrhythmias |  |  |  |
|  | Pathogenic test variants (TPR) |  |  | Benign test variants (TNR) |
|  | OMGL (N = 77) |  | HGMD (N = 138) | gnomAD (N = 1,237) |
| CardioBoost | **88.3\*\*\*** |  | **72.5\*\*\*** | **64.3\*\*\*** |
| M-CAP | 59.7 |  | 39.9 | 9.8 |
| REVEL | 68.8 |  | 52.9 | 2.8 |

### CardioBoost discriminates variants that are highly disease associated

Since benchmarking against a gold-standard test set may be susceptible to errors present in the benchmark data set, we employed two additional approaches to evaluate CardioBoost predictions directly against patient characteristics, to confirm biological and clinical relevance.

First, we directly assessed the strength of the association between the specified disease and rare variants stratified by the different tools. We compared the proportions of rare missense variants in a cohort of 6,327 genetically-characterised patients with HCM, from the SHaRe registry^20^, with 138,632 reference samples from gnomAD v2.0 (**Table 3**). We calculated the Odds Ratio (OR) of each sarcomere gene for all rare variants observed, and for variants stratified by CardioBoost, M-CAP, and REVEL after excluding variants seen in our training data.

**Table 3.** CardioBoost variant classification stratifies variants with increased disease Odds Ratio for sarcomere-encoding genes. Odd Ratios (ORs) and their confidence intervals were calculated for rare variants observed in sarcomere-encoding genes using SHaRe HCM cohorts and gnomAD. We compared the ORs for three groups of variants: (i) all rare variants, (ii) rare variants predicted pathogenic by CardioBoost (Pr ≥ 0.9, and excluding those seen in our training data), and (iii) rare variants predicted as benign by CardioBoost (Pr ≤ 0.1, and excluding those seen in our training data). The ORs of variants classified by M-CAP and REVEL were also calculated. For most of the sarcomere-encoding genes, variants classified as pathogenic by CardioBoost are enriched for disease-association, and those classified as benign are depleted, compared with unstratified rare missense variants.

| Gene symbol | all observed rare variants (95% CI) | CardioBoost pathogenic variants (95% CI) | CardioBoost benign variants (95% CI) | M-CAP pathogenic variants (95% CI) | M-CAP benign variants (95% CI) | REVEL pathogenic variants (95% CI) | REVEL benign variants (95% CI) |
| --- | --- | --- | --- | --- | --- | --- | --- |
| *MYH7* | 14.5<br>(14.4-14.6) | 14.7<br>(14.5-14.8) | 1.2<br>(0.7-1.7) | 14.8<br>(14.7-14.9) | -* | 15.9<br>(15.7-16.1) | -* |
| *TNNI3* | 12.6<br>(12.4-12.9) | 14.0<br>(13.1-14.8) | 3.3<br>(2.6-4.0) | 1.0<br>(1 -1.1) | 4.7<br>(3.7 – 5.8) | 12.1<br>(11-13.2) | 1.0<br>(0.9-1.1) |
| *TPM1* | 11.2<br>(10.7-11.7) | 33.7<br>(33.1 – 34.3) | 1.4<br>(0.4-2.4) | 1.0<br>(0.9 - 1.1) | 0.5<br>(0 - 2.5) | 38.9<br>(37-40.8) | -* |
| *ACTC1* | 11.2<br>(10.9-11.5) | 15.2<br>(14.6-15.8) | 1.0<br>(0.9-1.1) | 1.0<br>(0.9 - 1.1) | 1.0<br>(0.9 - 1.1) | 19.8<br>(19.1-20.6) | -* |
| *TNNT2* | 6.0<br>(5.8-6.2) | 17.7<br>(17.2-18.3) | 2.8<br>(2.2-3.4) | 1.0<br>(0.9 - 1.1) | 1.0<br>(0 - 3) | 25.8<br>(23.7-27.8) | 28.9<br>(27.1-30.6) |
| *MYBPC3* | 5.6<br>(5.5-5.6) | 55.1<br>(54.8-55.4) | 1.2<br>(0.9-1.4) | 1.0<br>(0.9-1.1) | 0.7<br>(0.2-1.2) | 12.8<br>(12.3-13.4) | 1.2<br>(0.8-1.6) |
| *MYL2* | 5.2<br>(5.0-5.5) | 3.8<br>(3.2-4.5) | 1.0<br>(0.9-1.0) | 1.0<br>(0.9-1.1) | 0.2<br>(0-2.2) | 1.7<br>(0.2-3.1) | 1.0<br>(0.9-1.1) |
| *MYL3* | 2.7<br>(2.3-3.0) | 7.9<br>(7.1-8.8) | 0.8<br>(0-1.7) | 1.0<br>(0.9-1.1) | 0.3<br>(0-2.3) | 19.4<br>(18.5-20.2) | -* |

For six out of eight CM-associated genes encoding sarcomere components (*TNNI3, TPM1, ACTC1, TNNT2, MYBPC3* and *MYL3*), the OR for variants prioritised by CardioBoost (i.e. predicted pathogenic with Pr ≥ 0.9) was significantly greater (*P-*value < 0.05) than the baseline OR (including all observed variants without discriminating pathogenic and benign variants), indicating that the tool is discriminating a set of pathogenic variants more strongly associated with the disease. Concordantly, variants in all the eight sarcomere genes predicted as benign have significantly decreased association with disease compared with the baseline OR (*P-*value < 0.05). By contrast, M-CAP or REVEL did not show any demonstrable difference in disease ORs between predicted pathogenic and predicted benign variants (**Table 3**).

### CardioBoost variant classification significantly associates with adverse clinical outcome

As a further assessment independent of gold-standard classification, we tested the association of variants stratified by CardioBoost with clinical outcomes in the same cohort of patients. Patients with HCM who carry known pathogenic variants in genes encoding sarcomeric proteins have been shown to follow an adverse clinical course compared with “genotype-negative” individuals (no rare pathogenic variant or VUS in a sarcomere-encoding gene, and no other pathogenic variant identified) ^20–22^, with a higher burden of adverse events. Patients carrying benign variants in HCM-associated genes would be expected to follow a similar trajectory to those genotype-negative patients.

We evaluated clinical outcomes in a subset of the SHaRe cohort comprising of 803 HCM patients each with a rare missense pathogenic variant or missense VUS in a sarcomere-encoding gene, and 1,927 genotype-negative HCM patients, after excluding all patients carrying variants that were seen in the CardioBoost training set. We compared event-free survival (i.e. age until the first occurrence of a composite adverse clinical outcome including heart failure events, arrhythmic events, stroke and death) of these patients, stratified by CardioBoost-predicted pathogenicity (the full definition of a composite adverse clinical outcome is described in **Supplementary Methods**).

CardioBoost classification stratifies novel variants with significantly different patient-survival curves (**Figure 3**). Patients carrying variants predicted as pathogenic (CardioBoost Pathogenic) were likely to have earlier onset and a higher adverse event rate than those without identified rare variants (CardioBoost Pathogenic vs Genotype negative: *P-*value < 2×10^−16^; Hazard Ratio (HR) = 1.9), or those with variants predicted to be benign (CardioBoost Pathogenic vs CardioBoost Benign: *P-*value = 0.03; HR = 1.7). The probability of developing the overall composite outcome by age 60 is 84% for CardioBoost Pathogenic patients, versus 60% for Genotype-negative patients. By contrast, groups stratified by M-CAP or REVEL variant classification did not show significantly different event-free survival time (M-CAP Pathogenic vs M-CAP Benign: *P*-value = 0.31; REVEL Pathogenic vs REVEL Benign: *P*-value = 0.30).

**Figure 3.**
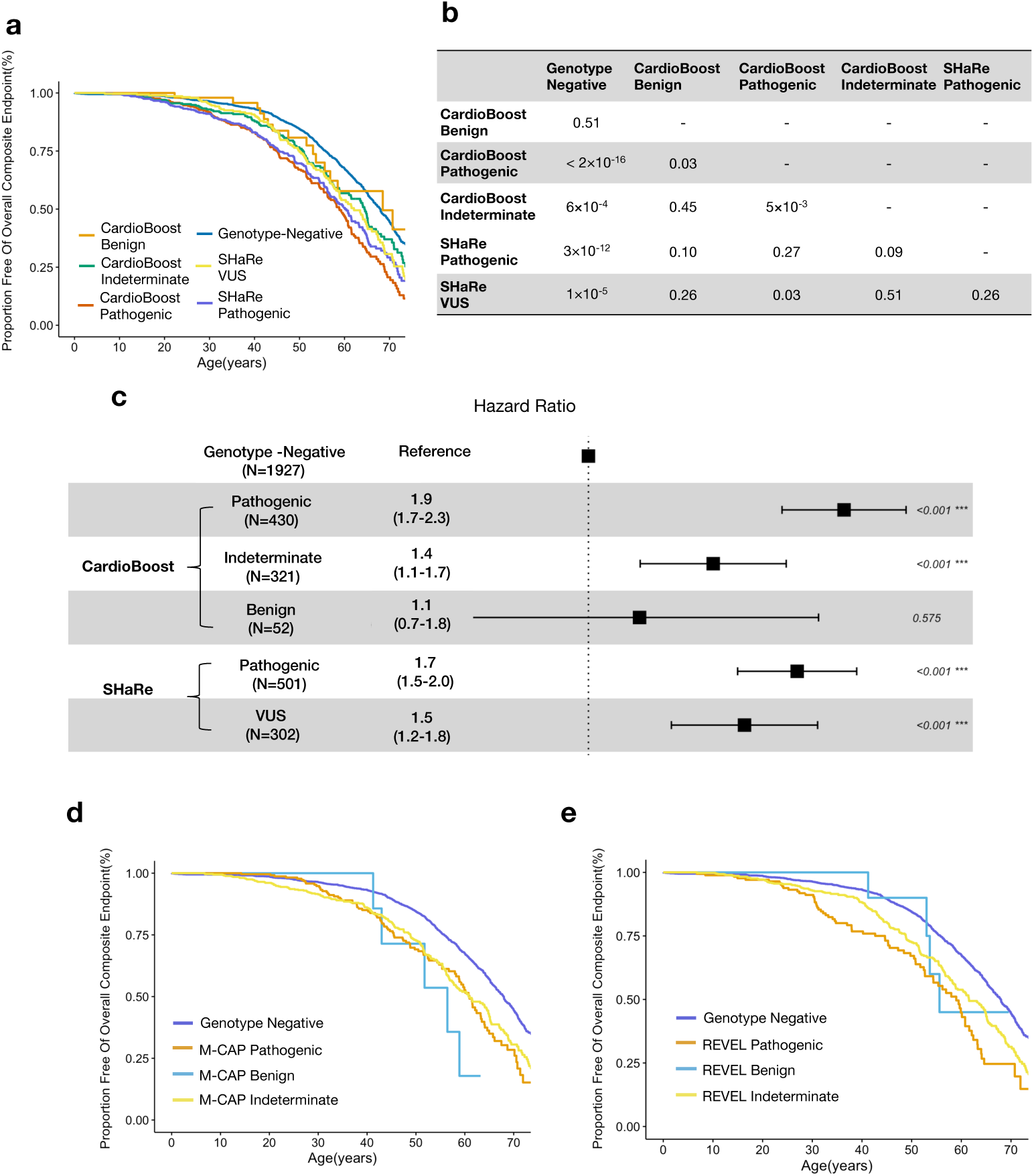
CardioBoost variant classification stratifies key clinical outcomes in patients with HCM. Clinical outcomes provide an opportunity to assess classifier performance independent of the labels used in the gold-standard training data. (**a**) Kaplan-Meier event-free survival curves are shown for patients in the SHaRe cardiomyopathy registry, stratified by genotype as interpreted by CardioBoost. The patients carrying variants seen in the CardioBoost training set were excluded in this analysis. Patients with pathogenic variants in sarcomere-encoding genes have more adverse clinical events compared with patients without sarcomere-encoding variants (“genotype-negative”), and compared with patients with sarcomere-encoding variants classified as benign. Survival curves stratified by variants as adjudicated by experts (marked in figure with prefix “SHaRe”) are shown for comparison. The composite endpoint comprised the first incidence of any component of the ventricular arrhythmic or heart failure composite endpoint, atrial fibrillation, stroke or death. (**b**) *P*-values of the log-rank test in the pairwise comparisons of Kaplan-Meier survival curves. (**c**) Forest plot displays the hazard ratio (with confidence interval) and *P*-value of tests comparing patients’ survival stratified by CardioBoost classification and SHaRe experts’ classification based on Cox proportional hazards models. Kaplan-Meier event-free survival curves for patients in the SHaRe cardiomyopathy registry, stratified by genotype as interpreted by M-CAP. The patients with variants predicted pathogenic by M-CAP did not have significantly different survival time compared to those with predicted benign variants (log-rank test *P*-value = 0.31). (**e**) Kaplan-Meier event-free survival curves for patients in the SHaRe cardiomyopathy registry, stratified by genotype as interpreted by REVEL. Patients with predicted pathogenic variants by REVEL did not have significantly different survival time compared to those with predicted benign variants (log-rank test *P*-value = 0.30).

## Discussion

Our results show that *in silico* prediction of variant pathogenicity for inherited cardiac conditions is improved within a disease-specific framework trained using expert-curated interpreted variants. This is demonstrated through improved classification performance, stronger disease-association, and significantly improved stratification of patient outcomes over published genome-wide variant classification tools.

There are several factors that may contribute to improved performance for a gene- and disease-specific classifier like CardioBoost over genome-wide tools. First, the use of disease-specific labels could decrease the false prediction of benign variants as pathogenic. A variant causative of one Mendelian dominant disorder may be benign with respect to a different disorder (associated with the same gene), if the conditions result from distinct molecular pathways. Since genome-wide tools are trained on universal labels (i.e. whether a variant ever causes any diseases), they would be expected to yield some false positive predictions in the context of specific diseases. Second, while the representative genome-wide tools M-CAP and REVEL are trained on variants from HGMD curated from literature, CardioBoost is trained on high-quality expert-curated variants, thus reducing label bias and increasing the prediction performances. Thirdly, as the genome-wide tools are trained across the genome, the learning function that maps the input features into the pathogenicity score is fitted using the training samples from all genes in the genome. However, different genes may have different mapping functions, for example related to different molecular mechanisms or the relevance of different features. Restricting to a set of well-defined disease-related genes may exclude influences from other unrelated genes.

We might expect a gene-disease-specific model would most accurately represent the genotype-phenotype relationship. However, there is a trade-off between the size of available training data and the specialization of prediction tasks. Here, CardioBoost groups together genes for two sets of closely related disorders, including three genes in which variants with different functional consequences lead to distinct phenotypes in our training set (i.e. *SCN5A, TNNI3, MYH7*). This is a potential limitation, since we hypothesise that distinct functional consequences might optimally be modelled separately. We explored alternative models for cardiomyopathy classifiers, for which our training data set is larger than for arrhythmias. Two disease-specific models (HCM-specific and DCM-specific) and three gene-syndrome-specific models (*MYH7*-HCM-specific, *MYH7*-DCM-specific, and *MYBPC3*-HCM-specific) with the largest training data size were built and compared (see **Supplementary Table 15**). None of the alternative models had comparable performance to the combined-cardiomyopathy model. We therefore conclude that given the current availability of training data, a cardiomyopathy-specific predictive model provides the best empirical balance between grouping variants with similar molecular or phenotypic effects and making use of relatively large training data set. It improves prediction both over genome-wide models that entirely ignore variants’ phenotypic effects, and over gene-disease-specific models for which there is insufficient training data. We therefore adopted the broadly disease-specific models as our final classifier, but anticipate that complete separation of distinct phenotypes may be advantageous when more training data becomes available in the future.

CardioBoost natively outputs a continuous probability of pathogenicity that is directly and intuitively interpretable. Users may therefore define their own confidence thresholds according to intended application. The posterior probability can also be updated by incorporating further evidence, such as linkage scores calculated from the evaluation of segregation in a family, to generate an updated posterior probability.

There are several further potential limitations and avenues for future refinement. First, we have only considered the prediction of pathogenicity for missense variants thus far. The inclusion of different classes of variants in disease-specific model is challenging since the available computational features or evidences for other types of variant are limited, and there is limited high-confidence training data for non-missense variants.

A second key limitation of CardioBoost is that it does not consider all relevant lines of evidence, and therefore it is not intended to serve as a tool for comprehensive assessment of variant pathogenicity. Some evidence types are limited by availability such as population allele frequency data and segregation data. Others could not be systematically included into a machine learning framework either because they are not well structured as in the case of functional data, de novo data and allelic data, or they are too sparse. For example, many variants lack experimental data, and the precise population allele frequency of many variants is unknown, though this implies significant rarity. In our training data, 45% of variants in cardiomyopathies and 44% of variants in arrhythmias were not seen in the gnomAD control population. Here, we do not model the imputation of absent allele frequencies in gnomAD for rare variants since the relation between variant pathogenicity and allele frequency scale beyond current observation is not clearly known.

For these reasons, while we show benefits of the proposed model for variant classification in known disease genes, and its superiority over existing genome-wide machine learning tools, we emphasize that CardioBoost is not intended for use as a standalone clinical decision tool, or as a replacement for the existing ACMG/AMP guidelines for clinical variant interpretation. Rather, in its current form it could provide a numerical value for evidence PP3 (“Multiple lines of computational evidence support a deleterious effect on the gene/gene product”) and BP4 (“Multiple lines of computational evidence suggest no impact on gene /gene product”) that is more reliable and accurate than existing genome-wide variant classifiers in the context of inherited cardiac conditions. We suggest that CardioBoost high-confidence classifications might appropriately activate PP3 (Pr>0.9) and BP4 (Pr<0.1). It is interpreted as the supporting evidence being activated with at least 90% confidence.

The widely-adopted ACMG/AMP framework is semi-quantitative, and the framework is largely internally consistent with a quantitative Bayesian framework^23^, but one limitation is that the weightings applied to different rules are not all evidence-based or proven to be mathematically well-calibrated. We do anticipate that, with more training data and robust validation, quantitative tools like CardioBoost will prove informative for variant interpretation, and will carry more weight in a quantitative decision framework than the current ACMG/AMP PP3 and BP4 rule affords.

As exemplified in two inherited cardiac conditions, we have substantiated that a disease-specific variant classifier improves the *in silico* prediction of variant pathogenicity over the best-performing genome-wide tools. We also demonstrate that development of a bioinformatic variant classifier represents a trade-off between biological specificity (i.e. a gene-disease-specific model) and practical availability of training data (i.e. a genome-wide model). For specific Mendelian disorders, it is important to understand the limitations of current genome-wide tools, and consider that a targeted gene-specific or disease-specific model may be advantageous given sufficient training data.

## Conclusions

We developed a machine-learning based variant classifier, CardioBoost, that is trained particularly on disease-specific variants to interpret rare missense variant pathogenicity on familial cardiomyopathies and inherited arrhythmias. In benchmarking with the existing genome-wide variant classification tools, CardioBoost significantly distinguishes more pathogenic and benign variants accurately with high confidence. Variants prioritised by CardioBoost with high confidence are also validated to be significantly associated disease state and predictive of patient survival in independent cohorts of cardiomyopathies. Our study also emphasizes the pitfalls of relying on genome-wide variant classification tools and the necessity to develop disease-specific variant classification tools to accurately interpret variant pathogenicity on specific phenotypes and diseases. We also highlight the need to evaluate variant classification tools in clinical settings including accuracies on high confidence classification thresholds equivalent to accepted certainty required for clinical decision making, variant association with disease and patients’ clinical outcomes. To support accurate variant interpretation in inherited cardiac conditions, we provide pre-computed pathogenicity scores for all possible rare missense variants in genes associated with inherited cardiomyopathies and arrhythmias (https://www.cardiodb.org/cardioboost/). The demonstrated development and evaluation framework could be applicable to develop accurate disease-specific variant classifiers and improve variant interpretation in a wide range of Mendelian disorders.

## List of Abbreviations

CM: (Inherited) Cardiomyopathy
FNR: False Negative Rate
FPR: False Positive Rate
gnomAD: Genome Aggregation Database release 2.0
HGMD: Human Genetics Mutation Database Pro version 201712
HR: Hazard Ratio
IAS: Inherited Arrhythmia Syndrome
ICC: Inherited Cardiac Condition
NPV: Negative Predictive Value
OMGL: Oxford Medical Genetics Laboratory
OR: Odds Ratio
PPV: Positive Predictive Value
PR-AUC: Area under the Precision-Recall Curve
Pr: Probability of pathogenicity
ROC-AUC: Area under the Receiver Operating Characteristic Curve
SHaRe: Sarcomeric Human Cardiomyopathy Registry version 2019Q3
TNR: True Negative Rate
TPR: True Positive Rate
VUS: Variant of Uncertain Significance
DM: Disease Mutation
ExAC: Exome Aggregation Consortium release 0.3
LMM: Laboratory of Molecular Medicine
MCC: Matthews Correlation Coefficient
RBH: Royal Brompton & Harefield Hospitals NHS Trust

## Ethics approval and consent to participate

Training and test data used in the development of the tool were either already in the public domain, or do not constitute personal data, or were obtained with patient consent and/or approval of the relevant research ethics committee or institutional review board.

## Availability of data and materials

The source code and data to reproduce our model development and validation analyses can be found on github at https://github.com/ImperialCardioGenetics/CardioBoost_manuscript. The pre-computed pathogenicity scores for all possible rare missense variants in genes associated with inherited cardiomyopathies and arrhythmias can be found at: https://www.cardiodb.org/cardioboost/.

## Authors’ contributions

XZ, LB and JSW designed the study and interpreted results. XZ developed the study and performed the analyses. RW, NW, RB, WM, AW, RG, NL, M.Ahmad, FM, AR, PT, EM, AdM, CJP, DPO’R, SAC, PJRB curated RBH data and provided feedback. M.Allouba, YA and MHY provided healthy volunteers data from Egypt Aswan Heart Centre and feedback. SMD, EA, SDC, MM, ACP, DJ, CYH, IO, GTG, JJ, CS and JI contributed the patient data from SHaRe cardiomyopathy registry. XZ, LB and JSW prepared the manuscript with input from co-authors. All authors reviewed and approved the final manuscript.

## Funding

The research was supported by the Wellcome Trust [107469/Z/15/Z; 200990/A/16/Z], British Heart Foundation [NH/17/1/32725; RE/18/4/34215], Medical Research Council (UK), National Institute for Health Research (NIHR) Royal Brompton Biomedical Research Unit, NIHR Imperial College Biomedical Research Centre, Science and Technology Development Fund (Egypt), Al-Alfi Foundation, Magdi Yacoub Heart Foundation, and the Alan Turing Institute under the Engineering and Physical Sciences Research Council grant [EP/N510129/1 to LB]. N.W. is supported by a Rosetrees and Stoneygate Imperial College Research Fellowship. JI is the recipient of a National Health and Medical Research Council (Australia) Career Development Fellowship (#1162929). C.S. is the recipient of a National Health and Medical Research Council (Australia) Practitioner Fellowship (#1154992).

## Competing interests

Professor Stuart Cook is a co-founder and director of Enleofen Bio PTE LTD, a company that develops anti-IL-11 therapeutics. Enleofen Bio had no involvement in this study. James Ware and Iacopo Olivotto have consulted for Myokardia, Inc. The ShaRe registry receives research support from MyoKardia. Myokardia had no involvement in this study.

## Acknowledgements

We thank Hugh Watkins and Kate Thomson (University of Oxford and Oxford Medical Genetics Laboratory) for making data available and for constructive discussion, and Mark Hazebroek (Maastricht University) for helpful feedback.

## References

1. Richards, S. et al. Standards and guidelines for the interpretation of sequence variants: a joint consensus recommendation of the American College of Medical Genetics and Genomics and the Association for Molecular Pathology. Genet Med 17, 405–423 (2015).

2. Kumar, P., Henikoff, S. & Ng, P. C. Predicting the effects of coding non-synonymous variants on protein function using the SIFT algorithm. Nat. Protoc. 4, 1073–1082 (2009).

3. Adzhubei, I., Jordan, D. M. & Sunyaev, S. R. Predicting functional effect of human missense mutations using PolyPhen-2. Curr. Protoc. Hum. Genet. 76, 7.20.1-7.20.41 (2013).

4. Schwarz, J. M., Cooper, D. N., Schuelke, M. & Seelow, D. Mutationtaster2: Mutation prediction for the deep-sequencing age. Nature Methods 11, 361–362 (2014).

5. Kircher, M. et al. A general framework for estimating the relative pathogenicity of human genetic variants. Nat. Genet. 46, 310–315 (2014).

6. Choi, Y., Sims, G. E., Murphy, S., Miller, J. R. & Chan, A. P. Predicting the Functional Effect of Amino Acid Substitutions and Indels. PLoS One 7, (2012).

7. Jagadeesh, K. A. et al. M-CAP eliminates a majority of variants of uncertain significance in clinical exomes at high sensitivity. Nat Genet 48, 1581–1586 (2016).

8. Ioannidis, N. M. et al. REVEL: An Ensemble Method for Predicting the Pathogenicity of Rare Missense Variants. Am. J. Hum. Genet. 99, 877–885 (2016).

9. Sundaram, L. et al. Predicting the clinical impact of human mutation with deep neural networks. Nat. Genet. 50, 1161–1170 (2018).

10. Anderson, D. & Lassmann, T. A phenotype centric benchmark of variant prioritisation tools. npj Genomic Med. 3, (2018).

11. Ruklisa, D., Ware, J. S., Walsh, R., Balding, D. J. & Cook, S. A. Bayesian models for syndrome- and gene-specific probabilities of novel variant pathogenicity. Genome Med. 7, 5 (2015).

12. Ackerman, J. P. et al. The Promise and Peril of Precision Medicine: Phenotyping Still Matters Most. Mayo Clin. Proc. 91, 1606–1616 (2016).

13. Hastie, T., Tibshirani, R. & Friedman, J. The elements of statistical learning: data mining, inference and prediction. (Springer, 2009).

14. Landrum, M. J. et al. ClinVar: improving access to variant interpretations and supporting evidence. Nucleic Acids Res. 46, D1062–D1067 (2018).

15. Cawley, G. C. & Talbot, N. L. C. On Over-fitting in Model Selection and Subsequent Selection Bias in Performance Evaluation. J. Mach. Learn. Res. (2010).

16. Saito, T. & Rehmsmeier, M. The precision-recall plot is more informative than the ROC plot when evaluating binary classifiers on imbalanced datasets. PLoS One 10, (2015).

17. Brier, G. W. VERIFICATION OF FORECASTS EXPRESSED IN TERMS OF PROBABILITY. Mon. Weather Rev. 78, 1–3 (1950).

18. DeLong, E. R., DeLong, D. M. & Clarke-Pearson, D. L. Comparing the Areas under Two or More Correlated Receiver Operating Characteristic Curves: A Nonparametric Approach. Biometrics 44, 837–845 (1988).

19. Stenson, P. D. et al. Human Gene Mutation Database (HGMD®): 2003 Update. Hum. Mutat. 21, 577–581 (2003).

20. Ho, C. Y. et al. Genotype and lifetime burden of disease in hypertrophic cardiomyopathy: insights from the Sarcomeric Human Cardiomyopathy Registry (SHaRe). Circulation 138, 1387–1398 (2018).

21. Lopes, L. R., Rahman, M. S. & Elliott, P. M. A systematic review and meta-analysis of genotype-phenotype associations in patients with hypertrophic cardiomyopathy caused by sarcomeric protein mutations. Heart 99, 1800–1811 (2013).

22. Ingles, J. et al. Nonfamilial Hypertrophic Cardiomyopathy. Circ. Cardiovasc. Genet. 10, (2017).

23. Tavtigian, S. V. et al. Modeling the ACMG/AMP variant classification guidelines as a Bayesian classification framework. Genet. Med. (2018). doi:10.1038/gim.2017.210

24. Ingles, J. et al. Evaluating the Clinical Validity of Hypertrophic Cardiomyopathy Genes. Circ. Genomic Precis. Med. 12, (2019).

