## Supplementary Methods for "Disease-specific variant pathogenicity prediction significantly improves variant interpretation in inherited cardiac conditions"

The data flow diagram from data collection, machine learning model training and testing is illustrated in **Figure 1**.

#### Primary training and test data collection

We consider rare missense variants whose allele frequency is less than 0.1%, using gnomAD (v2.0.1) as our reference population. The value at 0.1% is taken as a conservative maximum credible population allele frequency[1] across a range of inherited cardiac conditions, above which variants are unlikely to cause penetrant disease. The predicted molecular consequences of variants were annotated with Ensembl Variant Effect Predictor[2] (version 91.1 for hg19/GRCh37 human genome assembly) on canonical transcripts relevant to heart tissue (**Supplementary Table 1** and **Supplementary Table 2**).

Pathogenic variants in sixteen genes associated with cardiomyopathies (**Supplementary Table 1**) were collected from the targeted sequencing data of 9,007 patients with either HCM or DCM, recruited or referred for diagnostic sequencing at the Royal Brompton & Harefield Hospitals NHS Trust (RBH, UK), Oxford Medical Genetics Laboratories (OMGL, UK)[3], and the Partners Laboratory of Molecular Medicine (LMM, US)[4,5]. The pathogenic variants from RBH and OMGL were interpreted according to ACMG/AMP guidelines. The pathogenic variants from LMM were interpreted using equivalent previously-described clinical-grade variant classification criteria[4,5].

For inherited arrhythmia syndromes, pathogenic variants in seven genes **(Supplementary Table 2)** were extracted from the ClinVar database (ClinVar Full Release 201912), considering only variants with Pathogenic or Likely pathogenic classifications and no conflicting interpretations (Benign or Likely benign).

Rare benign variants for both conditions were collected from the targeted sequencing of 2,090 healthy volunteers: 921 from RBH, 423 from Aswan Heart Centre (Egypt)[6] and 746 from National Heart Centre (Singapore). These volunteers were confirmed to have no cardiac history, no family history of, or suggestive of, an inherited cardiac condition, and no evidence of cardiomyopathy or channelopathy on ECG or cardiac MRI. This cohort provides a lower disease prevalence than a general population (i.e. the prevalence of inherited cardiomyopathies and arrhythmias in a general population is estimated at ~0.75% by summing the combined prevalence of HCM, DCM, LQTS and Brugada syndrome[1]). Thus, the variants found in their disease panel genes could be considered as highly likely benign for inherited cardiac conditions, while acknowledging the potential for a low background error rate due to incomplete and age-related penetrance.

Three genes are each associated with two related disease phenotypes in the training & test data (*MYH7* and *TNNI3* with hypertrophic and dilated cardiomyopathies; *SCN5A* with two arrhythmia syndromes, LQT & BrS), with distinct variants causing each phenotype. For each of these genes variants were aggregated so that the model was trained to discriminate pathogenic for either condition vs. benign. The phenotype associated with variation in *PLN* does not fit neatly into the clinical definitions of either HCM or DCM[7], so the output of the model for *PLN* variants is interpreted as probability of variants causing intrinsic cardiomyopathy. For all other genes the model was exposed to variants associated with just one phenotype (HCM, DCM, BrS or LQT; **Supplementary Tables 1 & 2**).

**Additional replication test data collection**

To further validate CardioBoost performance on "unseen" data, we collected additional independent data sets which did not overlap with either the training data of CardioBoost, M-CAP and REVEL or the hold-out test data of CardioBoost.

For cardiomyopathies, these pathogenic test data sets are composed of 129

Pathogenic/Likely Pathogenic variants identified in HCM patients from the SHaRe Registry[8], 15 ClinVar (ClinVar Full Release 201912)[9] variants adjudicated as Pathogenic/Likely Pathogenic for cardiomyopathies with at least two-star review status, and 145 variants of the Disease Mutation (DM) class from HGMD Pro version 201712 after excluding those also seen in HGMD version 2015.2, since these variants were used in the training of M-CAP and REVEL. For arrhythmias, 77 variants reported to be Pathogenic/Likely Pathogenic by OMGL, and 138 variants of the DM class from HGMD Pro version 201712 were collected after excluding those seen in HGMD version 2015.2.

We expect most variants in disease-associated genes identified in gnomAD to be benign for inherited cardiac conditions since the prevalence of inherited cardiomyopathies and arrhythmias in gnomAD should not exceed those in a general population. Since ExAC[10] variants (ExAC version release 0.3, which represents a subset of gnomAD) were used to train M-CAP and REVEL explicitly, we curated a test set of 2,003 gnomAD variants in which the variants seen in ExAC were excluded. Similarly, for arrhythmias, 1,237 gnomAD variants were collected.

**Input variant features collection and pre-processing**

Feature collection. We combined both variant effect features collected from previous computational tools, and original newly-derived features.

There are two types of pre-existing computational tools for prediction of variant effect: (i) those that estimate the evolutionary conservation level of the genomic site or the variant itself; (ii) those that estimate the likelihood of variant pathogenicity combing both the conservation scores and biochemical properties of a variant. We used ANNOVAR[11] to collect features from published computational tools (**Supplementary Table 6**). Fourteen conservation or constraint scores of amino acid change were included from BLOSUM62[12], PAM250[12], Grantham Score[13], LRT[14], PhyloP[15], PhastCons[16], SIPHY[17], fitCons[18], GERP++[19], para_zscore[20] and misbadness[21]. To utilise the predictions of existing genome-wide tools, twenty pathogenicity scores were collected from SIFT[22], Polyphen2[23], MutationTaster[24], MutationAssessor[25], FATHMM[26], FATHMM-MKL[26], PROVEAN[27], VEST3[28], CADD[29], DANN[30], MetaSVM[31], MetaLR[31], Eigen[32], M-CAP[33], REVEL[34] and MPC[21].

To incorporate interspecies conservation maximally, we also derived new features measuring evolutionary conservation level from orthologous sequence alignments of disease genes. Using the multiple alignment of amino acid (AA) sequences of a set of species, for a given missense variant (with known site, reference AA and alternative AA) four types of features were extracted:

$$Ratio\ of\ Reference\ AA = \frac{\text{\#orthologs in the set that have the reference AA at that site}}{\text{\#orthologs in the set that have no gap at that site}}$$

$$Ratio\ of\ Alternative\ AA = \frac{\text{\#orthologs in the set that have the alternative AA at that site}}{\text{\#orthologs in the set that have no gap at that site}}$$

$$Ratio\ of\ No-Gap = \frac{\text{\#orthologs in the set that have no gap at that site}}{\text{\#orthologs in the set}}$$

$$Ratio\ of\ Orthologs = \frac{\text{\#orthologs in the set}}{\text{\#species in the set}}$$

We downloaded multiple sequence alignments of orthologous genes from the UCSC hg19 100-way Multiz alignment[35]. The above four scores were calculated for nine different sets of species: (1) all species included in the 100-way alignment; sets of species clade: (2) Primate (3) Euarchontoglires; (4) Laurasiatheria; (5) Afrotheria; (6) Mammal; (7) Aves; (8) Sarcopterygii and (9) Fish (For species in each clade subset see http://hgdownload.cse.ucsc.edu/goldenpath/hg19/multiz100way/).

We also derived region-level features from the AA alignment. $Mean\ Ratio\ of\ Reference\ AA$ measures the average ratio of $Ratio\ of\ Reference\ AA$ among the allele's 10 nearest neighbouring sites. Similarly, $Mean\ Ratio\ of\ No - Gap$ measures the average $Ratio\ of\ No - Gap$ among the allele's 10 nearest neighbouring sites.

Using the alignment of multiple nucleotide sequences, $Ratio\ of\ Reference\ Nucleotide$ and $Ratio\ of\ Alternative\ Nucleotide$ calculate the frequency of reference nucleotide and alternative nucleotide observed in all orthologs given there is no gap at this site respectively. Similarly, $Ratio\ of\ Reference\ Codon$ and $Ratio\ of\ Alternative\ Codon$ are derived as a measure of conservation at the codon level.

Missing features imputation. Variant pathogenicity scores derived from existing genome-wide classifiers and included as features in our model were not available for all variants considered. We estimated these missing values by using condition mean imputation. For test data, missing values were imputed by using the mean derived in the training data[36].

Features normalisation. In total, we collected 76 features per missense variant. After collecting all the features, we conducted a z-score normalisation on the features of the training data. The features in test data were also standardised using the means and standard variations of the training data.

**Defining high-confidence classification performance measures**

Existing machine learning variant classification tools adopted a single threshold to discriminate pathogenic and benign variants. However, the choice of this classification threshold is arbitrary and not consistent among different tools, for example M-CAP[33] made a binary classification using a threshold with 95% true positive rate (see the relevant discussion in **Supplementary**

**Methods**: Limitations in applying a high-sensitivity threshold for variant interpretation) and PolyPhen-2[23] made a ternary classification using two thresholds based on false positive rates.

This arbitrary choice of classification threshold might not be optimal in order to control Type I and Type II error for different applications. Moreover, the use of high-sensitivity threshold for variant classification is unlikely optimal for clinical interpretation of individual variants. Instead of using classification thresholds derived from a specific classification method/data set, here we adopt high-confidence classification definitions aligned with ACMG/AMP guideline recommendations for clinical practice[37]: the classification of variants into Likely Pathogenic/Pathogenic or Likely Benign/Benign is proposed to be with at least 90% classification certainty. In other words, variants with pathogenicity score equal to or larger than 0.9 would be classified as "pathogenic" and those with pathogenicity score equal to or smaller than 0.1 are classified as "benign". Variants with pathogenicity scores between 0.1 and 0.9 receive an indeterminate classification (variants of unknown significance) (**Figure 1b** and **Figure 1c**).

With the defined high-certainty classification thresholds, we derive the corresponding confusion matrix (**Figure 1c**) from which a series of measures of direct clinical relevance can be computed. We use TPR, the proportion of actual pathogenic variants predicted to be pathogenic, and PPV, the proportion of predicted pathogenic variants that are correctly classified, to evaluate the classifier's ability to classify pathogenic variants. TNR, the proportion of actual benign variants predicted to be benign and NPV, the proportion of predicted benign variants that are correctly classified are used to assess benign classifications correspondingly. Taking both cases together, the accuracy of high-confidence classifications measures the probability that a classification in the actionable range is correct. The proportion of clinically indeterminate classifications measures the probability of a variant not classified with clinical confidence. Formulae for each measure of clinical relevance we used are described in the below session.

### Calculation of high-confidence classification measures

**c**

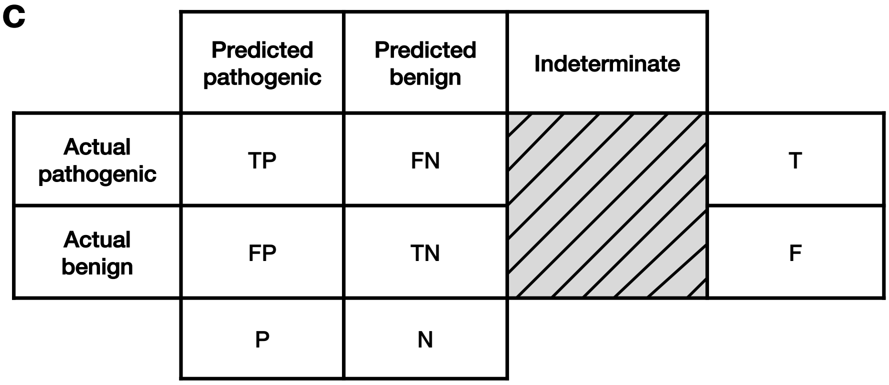


Based on the confusion matrix shown in **Figure 1c (shown above)**, we calculated the following ratios of clinical relevance in variant interpretation given *n* test variants

$$\mathrm{TPR} = \frac{\mathrm{TP}}{\mathrm{T}}$$

$$\mathrm{TNR} = \frac{\mathrm{TN}}{\mathrm{F}}$$

$$\mathrm{FPR} = \frac{\mathrm{FP}}{\mathrm{F}}$$

$$\mathrm{PPV} = \frac{\mathrm{TP}}{\mathrm{P}}$$

$$\mathrm{NPV} = \frac{\mathrm{TN}}{\mathrm{N}}$$

$$\mathrm{FNR} = \frac{\mathrm{FN}}{\mathrm{T}}$$

$$\text{Number of clinically relevant classifications} = \mathrm{P} + \mathrm{N}$$

$$\text{Number of indeterminate classifications} = n - (\mathrm{P} + \mathrm{N})$$

$$\text{Proportion of clinically relevant classifications} = \frac{\mathrm{P} + \mathrm{N}}{n}$$

$$\text{Accuracy of clinically relevant classifications} = \frac{\mathrm{TP} + \mathrm{TN}}{\mathrm{P} + \mathrm{N}}$$

$$\text{Overall accuracy} = \frac{\mathrm{TP} + \mathrm{TN}}{n}$$

$$\text{Proportion of indeterminate classifications} = \frac{n - (\mathrm{P} + \mathrm{N})}{n},$$

where T: Actual pathogenic, F: Actual benign, P: Predicted pathogenic (Pathogenicity $Pr \geq 0.9$), N: Predicted benign ($Pr \leq 0.1$), Indeterminate: $0.1 < Pr < 0.9$, TP: True Positive, TN: True Negative, FP: False Positive, FN: False Negative, T = TP + FN, F = FP + TN, P = TP + FP and N = FN + TN.

#### Machine learning model training and selection

The analyses were conducted using the R environment[23] and the package mlr[24]. We trained and tested representatives of each of the major classes of statistical and machine learning methods in order to obtain the best classification performances over our training data. Neural network methods were not included due to limited scope for interrogation and interpretation of feature weightings. Classification algorithms included in the analysis are: Classification and Regression Tree (CART)[40], K nearest neighbours (KNN)[41], Elastic Net Logistic Regression (GLMNET)[42], Support Vector Machine with Radial Basis Kernel Function (SVM-RBF)[43], Random Forest (RF)[44], Bayesian Additive Regression Trees (BART)[45], Adaptive Boosting (AdaBoost)[43], Gradient Boosting Tree (GBM)[43] and Extreme Gradient Boosting (XGBoost)[46].

To fine-tune hyper parameters for each model and identify the model with the best generalisation performance (i.e. best prediction performance on “unseen” data), we applied a nested cross-validation[47]. In this nested cross-validation, the inner-test set (also called “validation set” or “development set”) is used to choose the optimal set of hyperparameter for a given classification algorithm. After the classification algorithm is fitted on the inner loop data set, the outer test set is used to select the best tuned classification algorithm with respect to its performance on “unseen” test data. We used 5-fold cross-validation in the inner cross-validation loop and 10-fold in the outer cross-validation loop.

The selection of the best classification algorithm is not trivial. To this end, we pre-specified the following optimisation goals:

**Goal 1:** The optimal classifier outperforms genome-wide machine learning variant classification tools on overall classification measured using PR-AUC.

We consider the PR-AUC as a conventional threshold-independent performance measure. In the training process, PR-AUC is chosen as the objective measure in the inner loop for hyperparameter tuning, i.e. for each candidate classification algorithm considered, the hyperparameters that yield the highest PR-AUC are selected. Then the classification performance of each optimised algorithm is assessed using the outer CV loop.

**Goal 2:** The optimal classifier has the best Matthews Correlation Coefficient[48] (MCC) using the defined 90% high-confidence classification threshold.

Our aim is to find the optimal classifier that balances both Type I and Type II errors at the 90% high-confidence classification thresholds. When we apply the defined high-confidence classification above, variants are classified into one of three categories: pathogenic, benign and indeterminate. Since the most common application of a genetic diagnosis in cardiogenetic practice is familial evaluation and predictive testing, where management of negative and inconclusive genetic test results are equivalent[49], we group these variants together for the purposes of model selection, and focus on performance at the higher actionable threshold, comparing Likely Pathogenic/Pathogenic versus non-actionable indeterminate/Benign/Likely Benign.

We use the MCC, a measure of the correlation between observed and predicted binary classifications that is relatively robust in an imbalanced data set[50], defined as:

$$MCC = \frac{\mathrm{TP} \times \mathrm{TN} - \mathrm{FP} \times \mathrm{FN}}{\sqrt{(\mathrm{TP} + \mathrm{FP})(\mathrm{TP} + \mathrm{FN})(\mathrm{TN} + \mathrm{FP})(\mathrm{TN} + \mathrm{FN})}}.$$

A higher MCC reflects a stronger correlation between observed and predicted binary classification, indicative of performance at the ≥ 0.9 threshold most relevant in this context. Ideally, we would like to select a classifier that performs best on both goals. If there is more than one classifier satisfying both Goal 1 and Goal 2, we pre-specify selection of the models using Goal 2, given the most immediate relevance to this task.

The performance of each candidate machine learning algorithm and the benchmarking genome-wide variant classification tools (M-CAP and REVEL) in the nested cross-validation are shown in **Supplementary Table 7** and **Supplementary Table 8**. For cardiomyopathy variants, as shown in **Supplementary Table 7** the candidate algorithms that outperform M-CAP and REVEL on all standard classification measures to meet Goal 1 were GLMNET, CART, RF, BART, XGBoost, GBM, AdaBoost, KNN and SVM. Since AdaBoost had the highest MCC score to meet Goal 2, it was selected as the best model. Next the best hyperparameter set for AdaBoost (“loss=exponential” and “nu=0.207”) was selected using 5-fold cross validation on the whole cardiomyopathy variant training set. The selected model was trained on the whole training set to generate predictions on unseen data.

Similarly, for inherited arrhythmia syndrome variants, AdaBoost was selected as the best-performing candidate (hyperparameters “loss=exponential” and “nu=0.435”). The prediction model was then trained using the whole arrhythmia training set.

**Permutation significance test**

Given a performance measure, we used one-sided permutation test[51] to test whether an observed performance measure of one classifier was significantly better than that of the other classifier. The null hypothesis is that the two classifiers perform the same on this measure. The null distribution is estimated by randomly exchanging observations between the classifiers 10,000 times. Here, an observation represents a variant pathogenic probability predicted by a classifier. *P*-value is estimated as the number of times the permutated difference is larger than the observed difference.

**Replication without reliance on gold-standard**

To ensure robustness to misclassification in the “gold-standard” out-of-sample test data, we employed two orthogonal approaches to assess CardioBoost’s discrimination of pathogenic variants and benign variants. First, we compared the proportion of rare variants in individuals with and without disease, and stratified these variants using CardioBoost. We derived the odds ratio (OR), which provides an estimate of gene-disease association.

Second, we compared the survival outcomes of individuals with HCM, stratified by genotypes classified by CardioBoost. We applied CardioBoost to variants found in a cohort of 803 patients with HCM and a rare missense variant in one of eight HCM-associated genes, and compared survival with 1,927 genotype-negative HCM patients. We did not consider individuals carrying variants seen in our training data set. The “event-free survival” time (i.e. time until first major adverse clinical event) was analysed using Kaplan-Meier survival analysis and the Cox hazard-regression model.

**Survival analysis**

We collected genotype and clinical outcome data for patients with cardiomyopathy from the SHaRe HCM registry (data release 2019Q3).

We included patients with a diagnosis of HCM, at least 1 clinic visit and at least 1 assessment of left ventricular wall thickness, and only one missense variant in any of eight genes encoding sarcomere proteins (*MYBPC3*, *MYH7*, *TNNT2*, *TNNI3*, *TPM1*, *MYL2*, *MYL3*, and *ACTC1*). Variants jdentified in SHaRe were classified by SHaRe experts according to ACMG/AMP guidelines. Patients with potentially pathogenic variants in genes encoding non-sarcomere proteins (i.e. HCM genocopies) were excluded.

The primary outcome measure was a composite comprising the first occurrence of: sudden cardiac death, resuscitated cardiac arrest, appropriate implantable cardioverter-defibrillator therapy, cardiac transplantation, left ventricular assist device implantation, New York Heart Association class III-IV symptoms, all-cause mortality, atrial fibrillation, stroke, or death, as previously described[8].

Patients were censored either at date of first event, or at last follow-up clinical visit if event-free.

**Data leakage does not explain the superior performance of CardioBoost compared with existing tools**

Since CardioBoost training and test data may contain variants used as training data for published genome-wide classification tools whose pathogenicity scores were used as input features by CardioBoost, we also assessed whether using indirectly “seen" data would make CardioBoost overfit and outcompete existing genome-wide classifiers. In particular, we considered previously “seen” variants used in training M-CAP and REVEL. M-CAP was trained on variants of Disease Mutation (DM) Class from HGMD version 2015.2 and ExAC. REVEL was trained on variants of DMs from HGMD version 2015.2 and the Exome Sequencing Project (ESP), the Atherosclerosis Risk in Communities (ARIC) study and the 1000 Genomes Project (KGP) (ESP and KGP are contributing projects in ExAC). We extracted a set of “seen” variants from CardioBoost training data if they are ever seen in the DM Class of HGMD version 2015.2 and ExAC. The remaining variants in the training data constitute the set of purely “unseen” data. We investigated the impact of using “seen” data from two different viewpoints. One is whether “seen” variants have the same classification as those in our training data. In Cardiomyopathies 323 out of 440 training variants were seen before in HGMD or ExAC. For the DM variants reported in HGMD before, 53 out of 206 cases have an opposite classification as in our training data. In Arrhythmias, there are 253 out of 308 variants ever seen in HGMD or ExAC. Among the 170 DM variants reported in HGMD previously, 38 of them have opposite classification in our training data. This suggests that even if some variants were used in building previously genome-wide classifiers, their classifications are not necessarily correct and thus it makes the prediction tools less accurate. The second aspect is to assess whether our machine learning tool could still outcompete M-CAP and REVEL on completely “unseen” data. We compared the prediction performance of stratified hold-out test sets: purely “unseen” data and “seen” data (see **Table 1** and **Supplementary Table 9**) with the unstratified hold-out test set. The accuracy was used as an overall measure to compare the performance of each dataset. For cardiomyopathies and arrhythmias, the performances of three datasets were comparable and not significantly different. Overall, we found out the variants used in previous genome-wide tools were not necessarily accurately classified. Our machine learning tool did improve on cardiomyopathy- and arrhythmia-specific prediction both on “seen” and “unseen data” by leveraging over multiple diverse computational pieces of evidence.

**Limitations in applying a high-sensitivity threshold for variant interpretation**

In M-CAP, the authors defined a single low pathogenicity threshold as clinically relevant to predict pathogenic variants such that M-CAP could have 95% expected true positive rate (sensitivity). Given a data set, while using a low single classification threshold to increase TPR will decrease the number of false negative predictions, the binary classifier would tend to increase the number of false positive predictions (i.e., truly benign variants predicted to be pathogenic) as well. An ideal classification threshold would be the one that minimize the total sum of the cost of both errors. While one might prioritise sensitivity for variant prioritisation in some contexts, in the context of clinical variant interpretation, we suggest that the cost of a false positive prediction is at least equivalent to, and in most situations higher than, the cost of a false negative prediction. In neglecting to control the Type II error to have high true positive rate, there would be two negative consequences: (i) Low positive predictive value: this could be demonstrated as the negative correlation between the true positive rate and positive predictive value using the Precision-Recall Curve (**Figure 2a** and **Figure 2d**); (ii) High false positive rate: this is demonstrated as the positive correlation between the true positive rate and false positive rate (i.e., 1-TNR) (**Figure 2b** and **Figure 2e**). Even though the ACMG guidelines recommend not to use one computational tool as a sole evidence, but to consider the concordance of multiple computational tools for variant interpretation, the application of a computational tool of high TPR but low TNR or high FPR along with other computational tools would still make the clinical interpretation process rather difficult. For example, the pathogenic prediction of a computation tool for a truly benign variant is very likely to conflict with the correct prediction from the other computational tools or the other lines of evidence of pathogenicity. The contradictory evidence would increase the likelihood that the variant is classified as variant of uncertain significance (VUS).

**Calibration of PPV and NPV**

Given a new dataset or testing context, we could estimate the PPV and NPV of a classifier given the proportion of pathogenic variants amongst variants undergoing classification (Variant Proportion):

$$\text{Variant Proportion} = \frac{\text{Number of pathogenic variants}}{\text{Number of pathogenic variants} + \text{Number of benign variants}}$$

$$\text{PPV} = \frac{\text{TPR} \times \text{Variant Proportion}}{\text{TPR} \times \text{Variant Proportion} + \text{FPR} \times (1 - \text{Variant Proportion})}$$

$$\text{NPV} = \frac{\text{TNR} \times (1 - \text{Variant Proportion})}{\text{TNR} \times (1 - \text{Variant Proportion}) + \text{FNR} \times \text{Variant Proportion}}$$

where TPR: True Positive Rate and TNR: True Negative Rate as defined in (1) and (2), respectively.

**Estimating the proportion of pathogenic missense variants in a diagnostic series and a general population**

In order to estimate the PPV and NPV when applying CardioBoost in a diagnostic series and a general population, we first estimate the proportion of pathogenic missense variants of these two populations.

Since in variant interpretation, the limitation of false positive prediction is prioritised. Here we want to derive a reasonably conservative estimate of PPVs by assuming that pathogenic missense variants are penetrant and that the burden of rare missense variants in controls provides an estimate of the burden of rare benign missense variants in any population either cases or control. These assumptions would provide the lower bound of the proportion of pathogenic variants, which is the lower bound of PPV based on.

Based on the above assumptions, the proportion of rare pathogenic missense variants, for a given gene or a gene set, amongst variants identified in a group of patients with disorders could be approximated as:

$$\text{Variant proportion in a case series} = \frac{\text{Burden of pathogenic variants in cases}}{\text{Burden of rare variants in cases}}$$

$$\begin{aligned}&\text{Burden of pathogenic variants in cases}\\ &\qquad = \text{Burden of rare variants in cases} - \text{Burden of rare variants in control}\end{aligned}$$

Similarly, the proportion of rare pathogenic missense variants in a general population could be approximated as:

$$\begin{aligned}&\text{Variant proportion in a general population}\\ &\qquad = \frac{\text{Burden of pathogenic variants in a general population}}{\text{Burden of rare variants in a general population}}\end{aligned}$$

$$\begin{aligned}&\text{Burden of rare variants in a general population}\\ &\qquad = \text{Burden of pathogenic variants in a general population}\\ &\qquad + \text{Burden of benign variants in a general population}\end{aligned}$$

$$\begin{gathered}\text{Bruden of pathogenic variants in a general population} =\\ \text{Prevalence of disease } \times \text{Burden of pathogenic variants in cases} =\\ \text{Prevalence of disease } \times (\text{Burden of rare variants in cases} - \text{Burden of rare variants in control})\end{gathered}$$

$$\text{Burden of benign variants in a general population} = \text{Burden of rare variants in control}$$

For cardiomyopathies, here we consider both dilated cardiomyopathy (DCM) and hypertrophic cardiomyopathy (HCM). The disease prevalence for DCM is estimated as 1/250 and 1/500 for HCM[2]. Thus, adding the prevalence of two conditions, the disease prevalence for cardiomyopathies is

$$\frac{1}{250} + \frac{1}{500} \approx 0.006$$

Using cohort studies from OMGL and LMM[3], the burden of rare missense variants in cases is estimated at 27%. *PTPN11* (it was not sequenced in these cohorts and its contribution to cases is assumed to be marginal) was excluded in the analysis here. Using gnomAD[1] reference population as control, the burden of rare missense variants in control was estimated to be 11% by adding the allele frequencies of rare missense variants seen in gnomAD for all cardiomyopathies-related genes (excluding *PTPN11*).

Thus, the proportion of rare missense variants pathogenic to cardiomyopathies in a diagnostic series is estimated with ~ 60%. The proportion of rare missense variants pathogenic to cardiomyopathies in a general population is estimated as ~1%.

Likewise, the proportions of rare missense variants pathogenic to arrhythmias in a diagnostic series and in a general population are estimated as ~71% and ~0.4% respectively. The disease prevalence of arrhythmias in a general population is ~0.2% by adding the disease prevalence of Long QT syndrome (1/2000) and Brugada syndrome (1/1000). Since the arrhythmias-related genes are not widely assessed in large LQTS and Brugada cohort studies[4,5], here we could only consider four arrhythmias-associated genes *KCNE1, KCNH2*, *KCNQ1* and *SCN5A* here from the LQTS and Brugada cohort studies[4,5], which provides us a lower bound of exact variant proportion. The burden of rare missense variants in arrhythmias is estimated as 18%. From gnomAD database, we could estimate the burden of rare missense variants in control (only including *KCNE1, KCNH2*, *KCNQ1* and *SCN5A*) as 5%.

**References**


1. Whiffin, N. *et al.* Using high-resolution variant frequencies to empower clinical genome interpretation. *Genet. Med.* **19**, 1151–1158 (2017).
2. McLaren, W. *et al.* The Ensembl Variant Effect Predictor. *Genome Biol.* **17**, (2016).
3. Walsh, R. *et al.* Reassessment of Mendelian gene pathogenicity using 7,855 cardiomyopathy cases and 60,706 reference samples. *Genet Med* **19**, 192–203 (2017).

4. Pugh, T. J. *et al.* The landscape of genetic variation in dilated cardiomyopathy as surveyed by clinical DNA sequencing. *Genet. Med.* **16**, 601–608 (2014).
5. Alfares, A. A. *et al.* Results of clinical genetic testing of 2,912 probands with hypertrophic cardiomyopathy: Expanded panels offer limited additional sensitivity. *Genet. Med.* **17**, 880–888 (2015).
6. Aguib, Y. *et al.* Genomics of Egyptian Healthy Volunteers: The EHVol Study. *bioRxiv* (2019).
7. Ingles, J. *et al.* Evaluating the Clinical Validity of Hypertrophic Cardiomyopathy Genes. *Circ. Genomic Precis. Med.* **12**, (2019).
8. Ho, C. Y. *et al.* Genotype and lifetime burden of disease in hypertrophic cardiomyopathy: insights from the Sarcomeric Human Cardiomyopathy Registry (SHaRe). *Circulation* **138**, 1387–1398 (2018).
9. Landrum, M. J. *et al.* ClinVar: improving access to variant interpretations and supporting evidence. *Nucleic Acids Res.* **46**, D1062–D1067 (2018).
10. Lek, M. *et al.* Analysis of protein-coding genetic variation in 60,706 humans. *Nature* **536**, 285–291 (2016).
11. Wang, K., Li, M. & Hakonarson, H. ANNOVAR: functional annotation of genetic variants from high-throughput sequencing data. *Nucleic Acids Res.* **38**, e164–e164 (2010).
12. Henikoff, S. & Henikoff, J. G. Amino acid substitution matrices from protein blocks. *Proc. Natl. Acad. Sci.* **89**, 10915–10919 (1992).
13. Grantham, R. Amino Acid Difference Formula to Help Explain Protein Evolution. *Science.* **185**, 862–864 (1974).
14. Chun, S. & Fay, J. C. Identification of deleterious mutations within three human genomes. *Genome Res.* **19**, 1553–1561 (2009).
15. Siepel, A., Pollard, K. S. & Haussler, D. *New Methods for Detecting Lineage-Specific Selection*. *Research in Computational Molecular Biology* (Springer Berlin Heidelberg, 2006).
16. Siepel, A. *et al.* Evolutionarily conserved elements in vertebrate, insect, worm, and yeast genomes. *Genome Res.* **15**, 1034–1050 (2005).
17. Garber, M. *et al.* Identifying novel constrained elements by exploiting biased substitution patterns. *Bioinformatics* **25**, (2009).
18. Gulko, B., Hubisz, M. J., Gronau, I. & Siepel, A. A method for calculating probabilities of fitness consequences for point mutations across the human genome. *Nat. Genet.* **47**, 276–283 (2015).
19. Davydov, E. V. *et al.* Identifying a high fraction of the human genome to be under selective constraint using GERP++. *PLoS Comput. Biol.* **6**, (2010).
20. Lal, D. *et al.* Gene family information facilitates variant interpretation and identification of disease-associated genes. *bioRxiv* (2017). doi:10.1101/159780
21. Samocha, K. E. *et al.* Regional missense constraint improves variant deleteriousness prediction. *bioRxiv* (2017). doi:10.1101/148353
22. Kumar, P., Henikoff, S. & Ng, P. C. Predicting the effects of coding non-synonymous variants on protein function using the SIFT algorithm. *Nat. Protoc.* **4**, 1073–1082 (2009).
23. Adzhubei, I., Jordan, D. M. & Sunyaev, S. R. Predicting functional effect of human missense mutations using PolyPhen-2. *Curr. Protoc. Hum. Genet.* **76**, 7.20.1-7.20.41 (2013).
24. Schwarz, J. M., Cooper, D. N., Schuelke, M. & Seelow, D. Mutationtaster2: Mutation prediction for the deep-sequencing age. *Nature Methods* **11**, 361–362 (2014).
25. Reva, B., Antipin, Y. & Sander, C. Predicting the functional impact of protein mutations: Application to cancer genomics. *Nucleic Acids Res.* **39**, (2011).
26. Shihab, H. A. *et al.* Ranking non-synonymous single nucleotide polymorphisms based on disease concepts. *Hum. Genomics* **8**, (2014).
27. Choi, Y., Sims, G. E., Murphy, S., Miller, J. R. & Chan, A. P. Predicting the Functional Effect of Amino Acid Substitutions and Indels. *PLoS One* **7**, (2012).

28. Carter, H., Douville, C., Stenson, P. D., Cooper, D. N. & Karchin, R. Identifying Mendelian disease genes with the Variant Effect Scoring Tool. *BMC Genomics* **14**, S3 (2013).
29. Kircher, M. *et al.* A general framework for estimating the relative pathogenicity of human genetic variants. *Nat. Genet.* **46**, 310–315 (2014).
30. Quang, D., Chen, Y. & Xie, X. DANN: a deep learning approach for annotating the pathogenicity of genetic variants. *Bioinformatics* **31**, 761 (2014).
31. Dong, C. *et al.* Comparison and integration of deleteriousness prediction methods for nonsynonymous SNVs in whole exome sequencing studies. *Hum. Mol. Genet.* **24**, 2125–2137 (2015).
32. Ionita-Laza, I., McCallum, K., Xu, B. & Buxbaum, J. D. A spectral approach integrating functional genomic annotations for coding and noncoding variants. *Nat Genet* **48**, 214–220 (2016).
33. Jagadeesh, K. A. *et al.* M-CAP eliminates a majority of variants of uncertain significance in clinical exomes at high sensitivity. *Nat Genet* **48**, 1581–1586 (2016).
34. Ioannidis, N. M. *et al.* REVEL: An Ensemble Method for Predicting the Pathogenicity of Rare Missense Variants. *Am. J. Hum. Genet.* **99**, 877–885 (2016).
35. Kuhn, R. M., Haussler, D. & Kent, W. J. The UCSC genome browser and associated tools. *Brief. Bioinform.* **14**, 144–161 (2013).
36. Schafer, J. L. & Graham, J. W. Missing data: Our view of the state of the art. *Psychol. Methods* **7**, 147–177 (2002).
37. Richards, S. *et al.* Standards and guidelines for the interpretation of sequence variants: a joint consensus recommendation of the American College of Medical Genetics and Genomics and the Association for Molecular Pathology. *Genet Med* **17**, 405–423 (2015).
38. R Core Team. R: A Language and Environment for Statistical Computing. (2017).
39. Bischl, B. *et al.* mlr: Machine Learning in R. *J. Mach. Learn. Res.* **17**, 1–5 (2016).
40. Breiman, L., Friedman, J. H., Olshen, R. A. & Stone, C. J. *Classification and Regression Trees*. (Taylor & Francis, 1984).
41. Cover, T. & Hart, P. Nearest neighbor pattern classification. *IEEE Trans. Inf. Theory* **13**, 21–27 (1967).
42. Zou, H. & Hastie, T. Regularization and variable selection via the elastic-net. *J. R. Stat. Soc.* **67**, 301–320 (2005).
43. Hastie, T., Tibshirani, R. & Friedman, J. *The elements of statistical learning: data mining, inference and prediction*. (Springer, 2009).
44. Breiman, L. Random forests. *Mach. Learn.* **45**, 5–32 (2001).
45. Chipman, H. A., George, E. I. & McCulloch, R. E. BART: Bayesian additive regression trees. *Ann. Appl. Stat.* **6**, 266–298 (2012).
46. Chen, T. & Guestrin, C. *XGBoost : Reliable Large-scale Tree Boosting System. Proceedings of the 22nd ACM SIGKDD International Conference on Knowledge Discovery and Data Mining* (ACM, 2016).
47. Cawley, G. C. & Talbot, N. L. C. On Over-fitting in Model Selection and Subsequent Selection Bias in Performance Evaluation. *J. Mach. Learn. Res.* (2010).
48. Boughorbel, S., Jarray, F. & El-Anbari, M. Optimal classifier for imbalanced data using Matthews Correlation Coefficient metric. *PLoS One* **12**, (2017).
49. Cirino, A. L. *et al.* Role of genetic testing in inherited cardiovascular disease: A review. *JAMA Cardiol.* **2**, 1153–1160 (2017).
50. Chicco, D. Ten quick tips for machine learning in computational biology. *BioData Min.* **10**, (2017).
51. Good, P. I. *Resampling methods : a practical guide to data analysis*. (Birkhäuser, 2010).
52. Clark, T. G., Bradburn, M. J., Love, S. B. & Altman, D. G. Survival Analysis Part I: Basic concepts and first analyses. *British Journal of Cancer* **89**, 232–238 (2003).
