## Supplementary Tables for "Disease-specific variant pathogenicity prediction significantly improves variant interpretation in inherited cardiac conditions"

### List of Supplementary Tables

**Supplementary Table 1.** Cardiomyopathy-associated genes included in the study.

**Supplementary Table 2.** Arrhythmia-associated genes included in the study.

**Supplementary Table 3.** Data sets used for the development of CardioBoost.

**Supplementary Table 4.** The training data and hold-out test data grouped by gene used by CardioBoost for cardiomyopathies.

**Supplementary Table 5.** The training data and hold-out test data grouped by gene used by CardioBoost for arrhythmias.

**Supplementary Table 6.** Input variant features collected from existing computational tools.

**Supplementary Table 7.** Cross-validated out-of-sample performance for cardiomyopathy variant pathogenicity prediction.

**Supplementary Table 8.** Cross-validated out-of-sample performances for arrhythmia variant pathogenicity prediction.

**Supplementary Table 9.** Performance comparison on variants “unseen” and indirectly “seen” in the hold-out test data set for cardiomyopathy variant pathogenicity prediction.

**Supplementary Table 10**. Performance comparison on variants “unseen” and indirectly “seen” in the hold-out test data set for arrhythmia variant pathogenicity prediction.

**Supplementary Table 11.** CardioBoost outperforms existing genome-wide tools for the classification of hold-out test variants using 95%-certainty thresholds.

**Supplementary Table 12.** Comparison of classification performances on the hold-out test data set with minor allele frequency < 0.01%.

**Supplementary Table 13.** Evaluation of performances on additional test sets using 95%-certainty threshold.

**Supplementary Table 14.** Evaluation of performances on additional test sets with minor allele frequency < 0.01%.

**Supplementary Table 15.** Comparison of out-of-sample classification performances for alternative disease-specific classification models.

**Supplementary Table 1. Cardiomyopathy-associated genes included in the study.**

| Gene symbol | Phenotype | Ensemble gene ID | Ensemble transcript ID | Ensemble protein ID |
|---|---|---|---|---|
| *ACTC1* | HCM[1] | ENSG00000159251 | ENST00000290378 | ENSP00000290378 |
| *DES* | DCM[3] (syndromic) | ENSG00000175084 | ENST00000373960 | ENSP00000363071 |
| *GLA* | HCM[3] (syndromic) | ENSG00000102393 | ENST00000218516 | ENSP00000218516 |
| *LAMP2* | HCM[3] (syndromic) | ENSG00000005893 | ENST00000200639 | ENSP00000200639 |
| *LMNA* | DCM | ENSG00000160789 | ENST00000368300 | ENSP00000357283 |
| *MYBPC3* | HCM | ENSG00000134571 | ENST00000545968 | ENSP00000442795 |
| *MYH7* | HCM & DCM[1] | ENSG00000092054 | ENST00000355349 | ENSP00000347507 |
| *MYL2* | HCM | ENSG00000111245 | ENST00000228841 | ENSP00000228841 |
| *MYL3* | HCM | ENSG00000160808 | ENST00000395869 | ENSP00000379210 |
| *PLN* | Intrinsic CM[2] | ENSG00000198523 | ENST00000357525 | ENSP00000350132 |
| *PRKAG2* | HCM[3] (syndromic) | ENSG00000106617 | ENST00000287878 | ENSP00000287878 |
| *PTPN11* | HCM[3] (syndromic) | ENSG00000179295 | ENST00000351677 | ENSP00000340944 |
| *SCN5A* | DCM | ENSG00000183873 | ENST00000333535 | ENSP00000328968 |
| *TNNI3* | HCM & DCM[1] | ENSG00000129991 | ENST00000344887 | ENSP00000341838 |
| *TNNT2* | HCM[1] | ENSG00000118194 | ENST00000367318 | ENSP00000356287 |
| *TPM1* | HCM[1] | ENSG00000140416 | ENST00000403994 | ENSP00000385107 |

[1] While there are several genes in this table that have been associated with more than one type of cardiomyopathy, e.g. with different variants causing HCM and DCM, our training and test data included variants associated with just one type of cardiomyopathy for all genes except *MYH7 and TNNI3.* For *MYH7* and *TNNI3*, the output of CardioBoost should be interpreted as "probability of pathogenicity for HCM or DCM". For other genes associated with more than one subtype the classifier is trained for a particular disease only, and should be interpreted as such.

[2] The cardiomyopathic phenotype associated with variants in *PLN* does not fit neatly into the clinical definitions of HCM and DCM, so it has been classified under the broader umbrella of intrinsic cardiomyopathy[24].

[3] These conditions typically present with cardiomyopathy in the context of a broader syndromic phenotype, but may also present with isolated heart disease[24].

**Supplementary Table 2. Arrhythmia-associated genes included in the study.**

| Gene symbol | Phenotype | Ensemble gene ID | Ensemble transcript ID | Ensemble protein ID |
|---|---|---|---|---|
| *CACNA1C* | Timothy Syndrome (LQT) | ENSG00000151067 | ENST00000399655 | ENSP00000382563 |
| *CALM1* | LQT | ENSG00000198668 | ENST00000356978 | ENSP00000349467 |
| *CALM2* | LQT | ENSG00000143933 | ENST00000272298 | ENSP00000272298 |
| *CALM3* | LQT | ENSG00000160014 | ENST00000291295 | ENSP00000291295 |
| *KCNH2* | LQT | ENSG00000055118 | ENST00000262186 | ENSP00000262186 |
| *KCNQ1* | LQT | ENSG00000053918 | ENST00000155840 | ENSP00000155840 |
| *SCN5A* | LQT & BrS[1] | ENSG00000183873 | ENST00000333535 | ENSP00000328968 |

(LQT = Long QT syndrome; BrS = Brugada syndrome)

[1]For *SCN5A*, the output of CardioBoost should be interpreted as "probability of pathogenicity for LQT or BrS".

**Supplementary Table 3. Data sets used for the development of CardioBoost.** The number of missense variants in the training and hold-out test datasets is shown for two groups of inherited cardiac conditions.

| | Cardiomyopathies | | | Arrhythmias | | |
|---|---|---|---|---|---|---|
| | Pathogenic | Benign | Total | Pathogenic | Benign | Total |
| Training data set | 238 | 202 | 440 | 168 | 158 | 326 |
| Test data set | 118 | 100 | 218 | 84 | 79 | 163 |
| **Total** | **356** | **302** | **658** | **252** | **237** | **489** |

**Supplementary Table 4. The training data and hold-out test data grouped by gene used by CardioBoost for cardiomyopathies.** The number of missense variants in the training and hold-out test datasets is shown for each gene.

| | Training | Test |
|---|---|---|

| Gene symbol | Benign | Pathogenic | Benign | Pathogenic |
|---|---|---|---|---|
| *ACTC1* | 0 | 2 | 1 | 0 |
| *DES* | 13 | 3 | 4 | 0 |
| *GLA* | 5 | 5 | 3 | 3 |
| *LAMP2* | 5 | 2 | 1 | 0 |
| *LMNA* | 6 | 10 | 5 | 7 |
| *MYBPC3* | 47 | 19 | 27 | 14 |
| *MYH7* | 25 | 125 | 13 | 64 |
| *MYL2* | 1 | 11 | 1 | 1 |
| *MYL3* | 4 | 3 | 2 | 1 |
| *PLN* | 1 | 2 | 1 | 0 |
| *PRKAG2* | 14 | 2 | 7 | 2 |
| *PTPN11* | 8 | 1 | 2 | 1 |
| *SCN5A* | 55 | 2 | 27 | 0 |
| *TNNI3* | 6 | 23 | 4 | 8 |
| *TNNT2* | 8 | 14 | 2 | 8 |
| *TPM1* | 4 | 14 | 0 | 9 |
| **Total** | **202** | **238** | **100** | **118** |

**Supplementary Table 5. The training data and hold-out test data grouped by gene used by CardioBoost for arrhythmias.** The number of missense variants in the training and hold-out test datasets is shown for each gene.

| Gene symbol | Training | | Test | |
|---|---|---|---|---|
| | Benign | Pathogenic | Benign | Pathogenic |
| *CACNA1C* | 37 | 4 | 19 | 3 |
| *CALM1* | 0 | 5 | 0 | 1 |
| *CALM2* | 0 | 4 | 0 | 6 |
| *CALM3* | 0 | 3 | 0 | 0 |
| *KCNH2* | 33 | 54 | 19 | 22 |
| *KCNQ1* | 12 | 55 | 6 | 31 |
| *SCN5A* | 58 | 43 | 26 | 21 |
| **Total** | **140** | **168** | **70** | **84** |

**Supplementary Table 6. Input variant features collected from existing computational tools.**

| Features | Data type | Description |
|---|---|---|

| | | |
|---|---|---|
| Grantham score | Integer | Substitution matrix scoring the distance from one amino acid to the other |
| BLOSUM62 | Integer | |
| PAM250 | Integer | |
| SIFT | Float | Estimate intolerance to variation from closely-related species sequence alignment |
| Polyphen2 | Float x 2 | Machine learning method to predict functional effects using structural and sequence features |
| LRT_score | Float | The original LRT two-sided *P*-value |
| MutationTaster | Float | Bayes classifier used to predict pathogenicity of variants |
| MutationAssessor | Float | Predicts functional impact of amino acid substitutions |
| FATHMM | Float | HMM model to predict functional effects of variants |
| PROVEAN | Float | Predicts whether an amino acid substitution or indel has an impact on the biological function of a protein |
| VEST3 | Float | Machine learning method to predict variant functional effects |
| CADD | Float | SVM models to predict pathogenicity for coding and non-coding variants |
| DANN | Float | Scores whole-genome variants by training a deep neural network |
| FATHMM-MKL | Float | Machine learning method to predict variant functional effects |
| MetaSVM | Float | Machine learning method to predict SNVs functional effects |
| MetaLR | Float | Very similar to MetaSVM, but better interpretable |
| Eigen | Float x 2 | Unsupervised machine learning methods to predict function effects of coding and non-coding variants |
| M-CAP | Float | Gradient boosting tree to predict functional effects of missense variants |
| REVEL | Float | Random Forest to predict functional effects of missense variants |

| | | |
|---|---|---|
| GERP++ | Float | Identify constrained elements in multiple alignments |
| PhyloP | Float x 2 | Base pair level multi species conservation |
| Integrated_fitcons | Float | Estimate of fitness consequences |
| PhastCons | Float x 2 | Regional multi species conservation metric |
| SiPhy | Float | Detect bases under selection based on multiple alignments |
| paraZscore | Float | Estimate conservation across related proteins within-species from gene paralog |
| paraZscore_exist | Integer | Indicate whether the paraZscore of a missense variant is available |
| misbadness | Float | Measures the increased deleteriousness of amino acid substitutions when they occur in missense-constrained regions |
| misbadness_exist | Integer | Indicate whether the misbadness score of a variant is available |
| MPC | Float | Integrated score of misbadness, polyphen-2 and constraint |

**Supplementary Table 7. Cross-validated out-of-sample performance for cardiomyopathy variant pathogenicity prediction.** We compared nine classification algorithms including best-in-class representatives of all of the major families of machine learning algorithms. AdaBoost was selected with the best cross-validated out-of-sample performance. PR-AUC: Area under the Precision Recall Curve; ROC-AUC: Area under the Receiver Operating Curve; MCC: Mathew Correlation Coefficient.

| Method category | Algorithm | PR-AUC (%) | ROC-AUC (%) | Brier score | MCC |
|---|---|---|---|---|---|
| Regression | GLMNET | 90 | 88 | 0.15 | 0.10 |
| Tree-based | CART | 83 | 81 | 0.18 | 0.43 |
| | RF | 90 | 89 | 0.14 | 0.36 |
| | BART | 91 | 89 | 0.14 | 0.38 |
| Boosting-based | XGBoost | 90 | 87 | 0.15 | 0.51 |
| | GBM | 87 | 87 | 0.15 | 0.43 |
| | **Adaboost** | **90** | **88** | **0.14** | **0.58** |
| Other classification algorithms | KNN | 89 | 88 | 0.15 | 0.43 |
| | SVM-RBF | 89 | 87 | 0.14 | 0.36 |
| Existing genome-wide classification tools | M-CAP | 80 | 79 | 0.19 | 0.35 |
| | REVEL | 79 | 81 | 0.19 | 0.25 |

**Supplementary Table 8. Cross-validated out-of-sample performances for arrhythmia variant pathogenicity prediction.** We compared nine classification algorithms including best-in-class representatives of all of the major families of machine learning algorithms. AdaBoost was selected with the best cross-validated out-of-sample performance.

| Method category | Algorithm | PR-AUC (%) | ROC-AUC (%) | Brier score | MCC |
|---|---|---|---|---|---|
| Regression | GLMNET | 91 | 91 | 0.12 | 0.22 |
| Tree-based | CART | 82 | 86 | 0.14 | 0.56 |
| | RF | 93 | 92 | 0.10 | 0.45 |
| | BART | 93 | 92 | 0.11 | 0.43 |
| Boosting-based | XGBoost | 88 | 90 | 0.12 | 0.56 |
| | GBM | 87 | 89 | 0.12 | 0.60 |
| | **Adaboost** | **90** | **90** | **0.13** | **0.65** |
| Other classification algorithm | KNN | 92 | 91 | 0.12 | 0.45 |
| | SVM-RBF | 92 | 92 | 0.10 | 0.47 |
| Existing genome-wide classification tools | M-CAP | 81 | 85 | 0.16 | 0.38 |
| | REVEL | 89 | 90 | 0.17 | 0.59 |

**Supplementary Table 9. Performance comparison on variants “unseen” and indirectly “seen” in the hold-out test data set for cardiomyopathy variant pathogenicity prediction.** To assess whether bias is introduced in evaluating variants previously used in the training of M-CAP and REVEL, the performance of CardioBoost on wholly “unseen” data (not used in the training of M-CAP and REVEL), and indirectly “seen” data” (used in the training of M-CAP and

REVEL) were compared with M-CAP and REVEL. For each predictive performance measure (see **Supplementary Note** for details) the best algorithm is highlighted in bold.

| | "Unseen" data $N_{pathogenic}$ = 41 $N_{benign}$ = 24 | | | "Seen" data $N_{pathogenic}$ = 77 $N_{benign}$ = 76 | | |
|---|---|---|---|---|---|---|
| | CardioBoost (%) | M-CAP (%) | REVEL (%) | CardioBoost (%) | M-CAP (%) | REVEL (%) |
| PR-AUC | **90.2** | 80.2 | 73.8 | **91.8** | 78.6 | 76.7 |
| ROC-AUC | **86.3** | 71.1 | 70.2 | **92.1** | 79.8 | 81.9 |
| Brier Score | **13.4** | 21.5 | 19.5 | **11.8** | 19.0 | 19.2 |
| Overall Accuracy | **60.0** | 30.8 | 12.3 | **64.7** | 27.5 | 19.6 |
| Proportion of variants classified with high confidence | **69.2** | 40.0 | 20.0 | **70.6** | 31.4 | 22.9 |
| Accuracy of high-confidence classifications | **86.7** | 76.9 | 61.5 | **91.7** | 87.5 | 85.7 |
| Proportion of variants with indeterminate classifications | **30.8** | 60.0 | 80.0 | **29.4** | 68.6 | 77.1 |
| TPR | **70.7** | 43.9 | 19.5 | **68.8** | 40.3 | 32.5 |
| PPV | **82.9** | 75.0 | 61.5 | **88.3** | 86.1 | 83.3 |
| TNR | **41.7** | 8.3 | 0.0 | **60.5** | 14.5 | 6.6 |
| NPV | **100.0** | **100.0** | NA[1] | 95.8 | 91.7 | **100.0** |

1 No variants are classified as benign by REVEL.

**Supplementary Table 10. Performance comparison on variants "unseen" and indirectly "seen" in the hold-out test data set for arrhythmia variant pathogenicity prediction.** To assess whether bias is introduced in evaluating variants previously used in the training of M-CAP and REVEL, the performance of CardioBoost on entirely "unseen" data (not used in the training of M-CAP and REVEL), and indirectly "seen" data" (used in the training of M-CAP and

REVEL) were compared with M-CAP and REVEL. For each predictive performance measure (see **Supplementary Note** for details) the best algorithm is highlighted in bold.

| | "Unseen" data $N_{pathogenic}$ = 17 $N_{benign}$ = 18 | | | "Seen" data $N_{pathogenic}$ = 67 $N_{benign}$ = 52 | | |
|---|---|---|---|---|---|---|
| | CardioBoost (%) | M-CAP (%) | REVEL (%) | CardioBoost (%) | M-CAP (%) | REVEL (%) |
| PR-AUC | **94.4** | 82.2 | 87.1 | **96.8** | 88.6 | 93.1 |
| ROC-AUC | **94.1** | 85.6 | 86.3 | **95.0** | 84.6 | 92.6 |
| Brier Score | **12.2** | 15.9 | 20.6 | **9.3** | 17.4 | 16.2 |
| Overall Accuracy | **80.0** | 34.3 | 28.6 | **81.5** | 29.4 | 39.5 |
| Proportion of variants classified with high confidence | **88.6** | 40.0 | 34.3 | **88.2** | 31.9 | 42.0 |
| Accuracy of high-confidence classifications | **90.3** | 85.7 | 83.3 | 92.4 | 92.1 | **94.0** |
| Proportion indeterminate classifications | **11.4** | 60.0 | 65.7 | **11.8** | 68.1 | 58.0 |
| TPR | **88.2** | 70.6 | 58.8 | **82.1** | 43.3 | 67.2 |
| PPV | **88.2** | 85.7 | 83.3 | 91.7 | 93.5 | **93.8** |
| TNR | **72.2** | 0.0 | 0.0 | **80.8** | 11.5 | 3.8 |
| NPV | **92.9** | NA | NA | 93.3 | 85.7 | **100.0** |

**Supplementary Table 11. CardioBoost outperforms existing genome-wide classification tools for the classification of hold-out test variants using 95%-certainty thresholds.** While 90% is defined as a high-confidence threshold for clinical action in the ACMG/AMP guidelines, some may advocate a more stringent approach. We therefore assessed the performance of each tool using more stringent values for clinically relevant variant classification thresholds: high-confidence pathogenic (Pr ≥ 0.95), high-confidence benign (Pr ≤ 0.05), and indeterminate. For each predictive performance measure (see **Supplementary Note** for details) the best algorithm is highlighted in bold. Permutation tests were performed to evaluate whether the performance of CardioBoost was significantly different from the best value obtained by M-CAP or REVEL (significance levels: ****P*-value ≤ 0.001, ***P*-value ≤ 0.01, **P*-value ≤ 0.05).

| (%) | Cardiomyopathies | | | Arrhythmias | | |
|---|---|---|---|---|---|---|
| | CardioBoost | M-CAP | REVEL | CardioBoost | M-CAP | REVEL |
| Overall accuracy | **54.6*** ** | 16.5 | 7.3 | **78.6*** ** | 7.8 | 22.1 |
| Proportion of variants classified with high confidence | **60.1*** ** | 18.8 | 10.1 | **85.1*** ** | 8.4 | 23.4 |
| Accuracy of high confidence classifications | **90.8** | 87.8 | 72.7 | 92.4 | 92.3 | **94.4** |
| Proportion of variants with indeterminate classification | **39.9*** ** | 81.2 | 89.9 | **14.9*** ** | 91.6 | 76.6 |
| TPR | **62.7*** ** | 24.6 | 11.9 | **79.8*** ** | 11.9 | 39.3 |
| PPV | **87.1** | 85.3 | 70.0 | 91.8 | 90.9 | **93.9** |
| TNR | **45.0*** ** | 7.0 | 2.0 | **77.1*** ** | 2.9 | 1.4 |
| NPV | 97.8 | **100.0** | **100.0** | 93.1 | **100.0** | **100.0** |

**Supplementary Table 12. Comparison of classification performance on the hold-out test data set with minor allele frequency < 0.01%.** As novel pathogenic variants are more likely to be ultra-rare, CardioBoost was tested on the hold-out set of only ultra-rare variants and was confirmed to have comparable performance with that on rare variants. The performance of each tool is reported using the 90% high-confidence variant classification thresholds: high confidence pathogenic (Pr ≥ 0.90), high confidence benign (Pr ≤ 0.10), and indeterminate. For each predictive performance measure (see **Supplementary Note** for details) the best algorithm is highlighted in bold. Permutation tests were performed to evaluate whether the performance of CardioBoost was significantly different from the best value obtained by M-CAP or REVEL (significance levels: ***$P$-value ≤ 0.001, **$P$-value ≤ 0.01, *$P$-value ≤ 0.05).

| | Cardiomyopathies | | | Arrhythmias | | |
|---|---|---|---|---|---|---|
| | CardioBoost (%) | M-CAP (%) | REVEL (%) | CardioBoost (%) | M-CAP (%) | REVEL (%) |
| *Classification performance measures* | | | | | | |
| PR-AUC | **93*** | 85 | 81 | **97** | 90 | 95 |
| ROC-AUC | **91***** | 79 | 79 | **95** | 86 | 93 |
| Brier Score | **0.11*** | 0.18 | 0.17 | **0.09** | 0.15 | 0.14 |

| | | | | | | |
|---|---|---|---|---|---|---|
| *90% high-confidence classification performance measures* | | | | | | |
| Overall accuracy | **64.9*****| 30.9 | 19.7 | **83.6***** | 33.6 | 42.5 |
| Proportion of variants classified with high confidence | **71.3***** | 35.6 | 22.9 | **93.3***** | 93.8 | 95 |
| Accuracy of high confidence classifications | **91.0** | 86.6 | 86 | 93.3 | 93.8 | **95** |
| Proportion of variants with indeterminate classification | **28.7***** | 64.4 | 77.1 | **6.7***** | 65.2 | 53.3 |
| TPR | **70.1***** | 41.9 | 28.2 | **85.4***** | 50 | 67.1 |
| PPV | **89.1** | 86 | 84.6 | 94.6 | **95.3** | 94.8 |
| TNR | **56.3***** | 12.7 | 5.6 | **80.8***** | 7.7 | 3.8 |
| NPV | 95.2 | 90 | **100** | 91.3 | 80 | **100** |

**Supplementary Table 13. Evaluation of performances on additional test sets using 95%-certainty threshold.** CardioBoost performance was evaluated against five additional variant sets (SHaRe cardiomyopathy registry, ClinVar (two-star submissions), a UK regional genetic laboratory (Oxford Medical Genetics Laboratory – OMGL), the Human Gene Mutation Database – HGMD) and gnomAD after excluding the variants used to train CardioBoost, M-CAP and REVEL. The number of variants in each set is shown in brackets. The TPR is reported for pathogenic variant test sets (with threshold Pr ≥ 0.95), and the TNR for benign variant test sets (with threshold Pr ≤ 0.0.5). For each performance measure the best algorithm is highlighted in bold. Permutation tests were carried out to evaluate whether the performance of CardioBoost was significantly different from the best value obtained by M-CAP or REVEL (significance levels: ****P*-value ≤ 0.001, ***P*-value ≤ 0.01, **P*-value ≤ 0.05).

| (%) | Cardiomyopathies | | | |
|---|---|---|---|---|
| | Pathogenic test variants (TPR) | | | Benign test variants (TNR) |
| | SHaRe (N = 129) | ClinVar (N = 15) | HGMD (N = 145) | gnomAD (N = 2,003) |
| CardioBoost | **51.2*** ** | **60.0*** | **33.8*** ** | **44.2*** ** |
| M-CAP | 19.4 | 13.3 | 9.0 | 9.9 |
| REVEL | 6.2 | 6.7 | 6.9 | 2.6 |
| | Arrhythmias | | | |
| | Pathogenic test variants (TPR) | | | Benign test variants (TNR) |
| | OMGL (N = 77) | | HGMD (N = 138) | gnomAD (N = 1,237) |
| CardioBoost | **87.0*** ** | | **71.0*** ** | **61.3*** ** |
| M-CAP | 23.4 | | 18.8 | 4.3 |
| REVEL | 28.6 | | 23.9 | 1.2 |

**Supplementary Table 14. Evaluation of performances on additional test sets with minor allele frequency < 0.01%.**

(significance levels: ****P*-value ≤ 0.001, ***P*-value ≤ 0.01, **P*-value ≤ 0.05).

| (%) | Cardiomyopathies | | | |
|---|---|---|---|---|
| | Pathogenic test variants | | | Benign test variants |

| | (TPR) | | | (TNR) |
|---|---|---|---|---|
| | SHaRe (N = 129) | ClinVar (N = 14) | HGMD (N = 143) | gnomAD (N = 1,999) |
| CardioBoost | **62.0***** | **71.4*** | **42.0***** | **51.5***** |
| M-CAP | 37.2 | 42.9 | 22.4 | 20.3 |
| REVEL | 24.0 | 57.1 | 23.1 | 5.7 |
| | Arrhythmias | | | |
| | Pathogenic test variants (TPR) | | | Benign test variants (TNR) |
| | OMGL (N = 77) | | HGMD (N = 138) | gnomAD (N = 1,232) |
| CardioBoost | **88.3***** | | **72.5***** | **64.4***** |
| M-CAP | 59.7 | | 39.9 | 9.8 |
| REVEL | 68.8 | | 52.9 | 2.8 |

**Supplementary Table 15. Comparison of out-of-sample classification performances for alternative disease-specific classification tasks.** We explored alternative variant classification models as exemplified for cardiomyopathies with relatively larger size of training data: two syndrome-specific models (HCM-specific and DCM-specific) and three gene-syndrome-specific models (*MYH7*-HCM-specific, *MYH7*-DCM-specific and *MYBPC3*-HCM-specific). Here the broadly cardiomyopathies-specific model was chosen since none of the alternative models had comparable performances.

| Predictive task | Number of training variants | Precision-Recall AUC (%) |
|---|---|---|
| CM-specific | 440 | 91 |
| HCM-specific | 348 | 79 |
| DCM-specific | 309 | 48 |
| *MYH7*-HCM-specific | 152 | 87 |
| *MYH7*-DCM-specific | 152 | 35 |
| *MYBPC3*-HCM-specific | 106 | 76 |
